# Counteraction of HMGB1 at ss-dsDNA junctions maintains liquidity of protamine-DNA co-condensates

**DOI:** 10.64898/2026.02.24.707832

**Authors:** Vikhyaat Ahlawat, Divya Kota, Huan-Xiang Zhou

## Abstract

In the sperm nucleus, protamine replaces histones to mediate extreme DNA compaction. The histone-to-protamine transition involves the occurrence of double-strand breaks, and is facilitated by transition proteins including those containing high-mobility-group (HMG) boxes. Here we used optical tweezers and microscopy to study the actions of HMGB1 and protamine on DNA. Confocal scans of GFP-HMGB1 on overstretched λ-DNA show 2-3 foci that spread on the DNA upon retraction. Spreading of foci coincides with reannealing of ssDNA tracks, confirming their localization at ss-dsDNA junctions. Whereas the force–extension curves of protamine-bound λ-DNA show tangles that withstand forces > 60 pN, premixing protamine with HMGB1 produces only bends and bridges (∼ 20 pN). The counteraction of HMGB1 involves its acidic C-terminal tail, as HMGB1-ΔC fails to prevent tangle formation. In line with these single-molecule results, brightfield and confocal imaging shows that HMGB1 converts protamine–dsDNA aggregates into liquid droplets whereas HMGB1-ΔC fails to do so. Together, these observations support our hypothesis that chromatin-associated proteins like HMGB1 help maintain early protamine-mediated DNA condensates in a liquid state, enabling the recruitment of the repair machinery to restore the duplex structure.

## Introduction

In the sperm nucleus, protamines replace histones to achieve extreme chromatin compaction.^1–4^ In mammals and fish, protamines are small Arg-rich disordered proteins. In flies, protamine-like proteins including protamine A, protamine B, and Mst77F^5^ belong to a group of proteins known as sperm nuclear basic proteins (SNBPs; Table S1). SNBPs each contain one or two high-mobility group (HMG) boxes and Arg/Lys-rich motifs.^6^ DNA supercoiling builds up as histones are removed; topoisomerase IIβ creates double-strand breaks to relieve supercoiling.^7–9^ Transition proteins (Arg/Lys-rich disordered proteins in mammals) are then transiently expressed as protamines start to appear.^10–12^ In flies, a subset of SNBPs, including Tpl94D,^8^ acts as transition proteins. Other chromatin-associated proteins may also play a role. For example, the prototypical HMG box-containing proteins, HMGB1 and HMGB2, are transiently expressed in the mouse testis; HMGB2 deletion caused defective spermiogenesis.^13, 14^ In the meantime, double-strand breaks trigger the activation of PARP1 and PARP2,^15^ which may then recruit repair factors to restore the duplex structure of DNA. Finally protamine condenses double-stranded DNA (dsDNA) into hyperstable “tangles”,^16^ leading to global transcription silencing.

Fluorescence imaging demonstrated highly efficient condensation of a single λ-DNA concatemer by salmon protamine, which contains 21 Arg residues in its 33-residue sequence.^17^ The same condensation rate was achieved by a six-Arg peptide at a higher concentration (160 vs 2 µM). Similar observations were obtained with bull protamine 1, which contains 26 Arg residues in its 51-residue sequence.^18^ Arg-to-Lys substitutions lowered the condensation rate and raised the decondensation rate, resulting in a significant decrease in effective DNA binding affinity. A Lys-to-Ala substitution in mouse protamine 1 produced a similar effect in DNA curtain assays.^19^ Early atomic force microscopy (AFM) and transmission electron microscopy of sperm chromatin suggested toroidal structures.^20^ Recent AFM studies of single DNA chains condensed by salmon protamine found a variety of structures, including stacks of loops,^21^ coils, rods, and toroids.^22^ Optical tweezers (OT)-based single-molecule force spectroscopy showed that salmon protamine condensed single dsDNA chains into tangles that withstand forces strong enough (∼60 pN) to separate the two strands of DNA, along with bends and loops that unravel at 10-40 pN forces. As their strength far exceeds the stall force (7.5-25 pN) of RNA polymerases,^23–25^ protamine-mediated tangles provide a mechanism for global transcription silencing. All-atom molecular dynamics simulations revealed that Arg sidechains not only form salt bridges with the phosphate backbone of DNA but also bifurcated hydrogen bonds (“wedges”) with nucleobases from both strands simultaneously, potentially explaining the hyper-stability of tangles.^16^ In bulk experiments, protamine condensed dsDNA into solid-like aggregates.^16, 26, 27^

It can be argued that, early in spermiogenesis when protamines are first expressed, chromatin needs to be maintained in a liquid state in order for the recruitment of repair factors. Indeed, premature synthesis of protamine 1 in mouse spermatids resulted in precocious condensation of nuclear DNA.^28^ In addition, posttranslational modifications of protamines may regulate the extent of chromatin condensation during spermiogenesis^29^ and embryogenesis.^26^ Interestingly, co-condensation of protamine with single-stranded DNA (ssDNA) produced liquid droplets^27, 30^ as opposed to solid-like aggregates with dsDNA. Correspondingly, single-molecule force spectroscopy of protamine-condensed ssDNA showed formation of bridges, which was characterized by a sawtooth pattern and ruptured at ∼20 pN force.^30^ Moreover, the magnitude of the bridge-rupture force can be tuned by the ssDNA-to-dsDNA ratio. These observations led us to suggest that the presence of ssDNA tracks during histone-to-protamine transition might contribute to maintaining the liquidity of DNA condensates.

HMGB1 can potentially serve as a model protein for increasing the liquidity of protamine-dsDNA co-condensates. As noted above, this protein and its close homolog HMGB2 are transiently expressed during spermiogenesis of mice.^13^ Also, SNBPs that mediate spermiogenesis of flies all contain HMG boxes (Table S1). Importantly, in somatic cells, HMGB1 counters linker histone H1 to loosen chromatin and increase the accessibility of nucleosomal DNA, for transcription and DNA repair.^31–37^ HMGB1 comprises two tandem HMG boxes followed by a 30-residue track of Asp and Glu. The acidic tail is required for HMGB1’s stimulation of transcription.^38^ The HMG boxes bind to the minor groove and bend dsDNA.^39–41^ Correspondingly, HMGB1 preferentially binds to distorted or damaged sites on DNA.^42–44^ The acidic tail competes with DNA for binding to the HMG boxes.^45–47^ A tail-deletion variant, HMGB1-ΔC (or ΔC for short), increased both the flexibility and contour length of λ-DNA in a single-molecule study.^48^ Compared to HMGB1-ΔC, the full-length protein has a substantially lower affinity for binary complex formation with dsDNA (Kd ∼2 nM^48^ versus 100 nM^37^) and a ∼10-fold faster dissociation rate.^47^ HMGB1 also uses its acidic tail to interact with the basic tail of histone H1.^34^ This interaction may help HMGB1 compete against H1 for DNA binding. Indeed, live-cell fluorescence recovery after photobleaching (FRAP) showed that exogenous HMGB1 decreased the residence time of H1 in chromatin.^33^ Recent cryo-electron microscopy and biochemical studies suggest that HMGB1 counters the effects of H1 by forming a ternary complex with DNA, where the interaction of HMGB1’s acidic tail with the basic tail of H1 loosens the latter’s binding with DNA.^37^ With the help of PEG as a crowding agent, the interactions of the acidic tail and the HMG boxes on different HMGB1 molecules drive homotypic droplet formation.^49^

Here we combined single-molecule studies and bulk experiments to characterize the counteraction of HMGB1 against salmon protamine to maintain the liquidity of DNA condensates. On overstretched λ-DNA, HMGB1 co-condenses with peeled-off ssDNA tracks and thereby forms foci at ss-dsDNA junctions, similar to protamine,^16^ histone H1,^50^ and prion-like protein FUS.^51^ However, unlike these proteins, HMGB1 foci spread to the adjacent unpeeled ssDNA sections and promote strand reannealing. In addition, while protamine produces a large condensing force (∼10 pN) in λ-DNA held at low stretch and induces the formation of hyperstable tangles, HMGB1 does not condense relaxed λ-DNA and merely increases its flexibility. When HMGB1 is present with protamine, it converts tangles into bridges that rupture at ∼25 pN; this conversion requires the acidic tail. In line with these single-molecule observations, HMGB1 converts protamine-dsDNA aggregates into liquid droplets. Together, these results suggest that HMGB1, via its acidic tail, can peel off protamine away from DNA and maintain condensate liquidity, with profound implications for spermiogenesis.

## Results

To mimic strand breaks during the histone-to-protamine transition in spermiogenesis, we overstretched λ-DNA by dual-trap OT to unpeel ssDNA tracks from nicks. We then conducted correlative force-fluorescence measurements of the λ-DNA tethers in the presence of HMGB1 alone or both HMGB1 and protamine, and collected additional force-extension curves without doping of the GFP-HMGB1 fusion protein. These single-molecule studies were complemented by bulk experiments, including brightfield and confocal imaging of protein–DNA mixtures for condensation propensities and material states and FRAP and OT-directed fusion for condensate dynamics.

### HMGB1-mediated condensates at ss-dsDNA junctions spread over the DNA surface and promoted strand reannealing

We first recapitulate the essential properties of protamine-mediated condensates on DNA observed previously,^16, 30^ in order to draw contrast with the behaviors of HMGB1-mediated condensates. When overstretched λ-DNA tethers (extension ∼24 µm) were brought into the protamine channel (with SYTOX Orange for staining) and retracted, foci emerged at ss-dsDNA junctions and grew in size, as captured by kymographs (Figure 1a) and 2D scans (Figure 1b). The growth in size, quantified by fluorescence intensity (Figure 1c), results from dsDNA being drawn into condensates. Force-extension curves of initially overstretched λ-DNA in the protamine channel (without the dye) provided additional insight (Figure 1d). The retraction curve lagged the counterpart of intact DNA in buffer (“naked” DNA), indicating incomplete reannealing of ssDNA tracks. Moreover, a characteristic dip at ∼ 11 µm marked a transition from ssDNA-dominated to dsDNA-dominated condensates; the final baseline at ∼ 10 pN (instead of going to 0 pN) signified condensation. The subsequent stretching curve exhibited a much earlier rise to strand separation than naked DNA, signifying the formation of tangles (hyperstable nested loops crosslinked by protamine).

**Figure 1.**
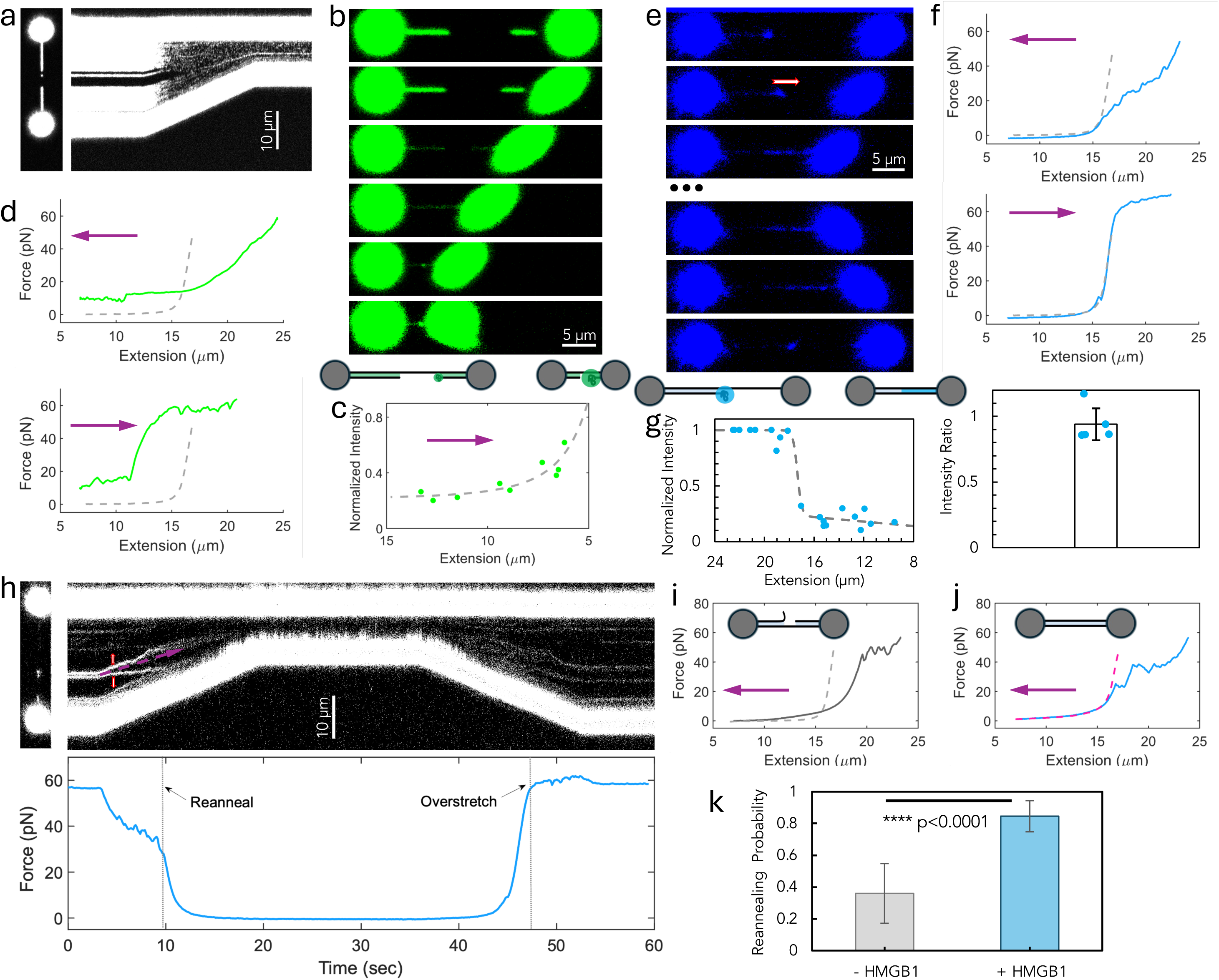
Contrasting behaviors of protamine- and HMGB1-mediated condensates at ss-dsDNA junctions. (**a**) A kymograph showing the emergence of a focus from an ss-dsDNA junction and subsequent growth in size as an λ-DNA tether (initially overstretched to ∼ 24 µm in the buffer channel) was retracted to 7 µm in the protamine channel (10 µM protamine plus 0.5 µM SYTOX Orange for staining). Fluorescence was rendered black and white. A 2D scan of the λ-DNA at its initial overstretched state is also included. (**b**) 2D scans of initially overstretched λ-DNA undergoing retraction in the protamine channel. The cartoon at the bottom depicts condensate formation and growth at an ss-dsDNA junction. (**c**) Growth in fluorescence intensity inside foci with increasing λ-DNA retraction. Fluorescence intensities were measured over a given focus over successive scans and normalized to the value on the overstretched λ-DNA in the first scan; results from three λ-DNA tethers were pooled. The curve is a fit to equation [1]. (**d**) Force-extension curves of an initially overstretched λ-DNA tether in the protamine channel, during retraction (top) and subsequent stretch (bottom). Here and in subsequent figures displaying force-extension curves, the stretching/retraction curve of naked DNA is shown as gray dash for reference; leftward and rightward arrows mean retraction and stretching, respectively. (**e**) 2D scans of λ-DNA during its retraction and subsequent stretch in the presence of HMGB1. The initially overstretched λ-DNA tether was soaked in the HMGB1 channel (8 µM HMGB1 and 0.8 µM GFP-HMGB1) and moved just outside the channel (to reduce the GFP-HMGB1 background) for scanning (see Figure S2a). A red arrow in the first frame indicates that the focus migrated in the opposite direction of retraction. Six frames in the middle of the sequence are skipped (indicated by ellipses). The cartoon at the bottom depicts condensate spreading and strand reannealing. (**f**) Force-extension curves corresponding to the retraction and stretching in (**e**). Top: merge of the retraction curve with the naked-DNA reference indicates completion of strand reannealing. Bottom: overlap of the stretching curve with the naked-DNA reference confirms complete strand reannealing. (**g**) Spreading of HMGB1-mediated condensates along λ-DNA. Left: change in fluorescence intensity inside foci as the λ-DNA tether was retracted, showing a switch-like transition indicating condensate spreading over the DNA surface. After spreading, the fluorescence intensity was the mean of the values measured at three positions inside the spread. All intensities were normalized by the total intensity inside all foci in the first scan; results from five λ-DNA tethers were pooled. The curve is a fit to equation [2]. Right: similar to the left panel, but the fluorescence intensity in the entire spread was integrated for the scan where spreading first occurred. The error bar displays the standard deviation among five tethers. (**h**) Top: a kymograph capturing the same process as by 2D scans of (**e**). A slanted arrow indicates the would-be direction of a static point during retraction; vertical arrows indicate the migration directions of two nearby foci. Bottom: force trace, showing that condensate spreading and reemergence coincided with strand reannealing and separation, respectively. (**i**) A retraction curve of an overstretched tether in buffer showing residual strand separation, as indicated by a lag from the counterpart (gray dash) of intact DNA. (**j**) Similar to (**i**) but retraction occurred in the presence of 5 µM HMGB1, showing complete reannealing when the extension decreased to 16 µm. The retraction curve of intact DNA in the presence of 5 µM HMGB1 is displayed in magenta dash for reference. (**k**) Reannealing probabilities of overstretched λ-DNA without and with 5 µM HMGB1. Error bars display standard deviations among 52 (or 25) stretch-retract cycles of 20 (or 13) tethers with (without) HMGB1.

A single-molecule study of McCauley et al.^48^ showed that HMGB1-ΔC at nM concentrations increased both the flexibility and contour length of λ-DNA. Given the much lower affinity of full-length HMGB1 relative to HMGB1-ΔC,^37, 48^ here we investigated the effects on DNA by HMGB1 and HMGB1-ΔC at µM concentrations. Consistent with McCauley et al., force-extension curves in the presence of µM ΔC revealed increases in both the flexibility and contour length of λ-DNA; in contrast, µM HMGB1 increased the flexibility but not the contour length (Figure S1).

When an overstretched λ-DNA tether (extension ∼23 µm) was soaked in 8 µM HMGB1 (doped with 0.8 µM GFP-HMGB1 for fluorescence) and then retracted in buffer (Figure S2a), an HMGB1 focus emerged as a thin coating covered the presumed dsDNA section of the tether (Figure 1e first scan). It looked as if the focus was formed with peeled-off ssDNA and at the ss-dsDNA junction, similar to what was observed with protamine during initial retraction as well as histone H1^50^ and FUS^51^. However, in departure from these proteins, as retraction proceeded, the HMGB1 focus migrated toward the adjacent (presumed) ssDNA section (Figure 1e second scan) and then spread to the entire ssDNA section (Figure 1e third scan). The corresponding force-extension curve (Figure 1f top) showed that ssDNA tracks completed reannealing at this extension. The coincidence of condensate spreading and strand reannealing provided additional evidence that the initial focus was located at the ss-dsDNA junction. Also of note was the final merging of the retraction curve with the baseline, indicating the absence of dsDNA condensation. In the subsequent stretch, the force-extension curve nearly overlapped with the naked-DNA reference, confirming complete strand reannealing (Figure 1f bottom). With a further increase in extension to the overstretched regime, the spreading reverted into a focus (Figure 1e, last scan). The coincidence of condensate spreading and strand reannealing was reproducible over multiple λ-DNA tethers (Figure 1g left). The total fluorescence intensity in the entire spread at first occurrence (third or second scan) was nearly the same as that in the initial foci (typically two or three per tether; Figure 1g right), reflecting the fact that the scans took place in buffer and hence HMGB1 could not be replenished. The intensity faded over time as HMGB1 dissociated into the surrounding buffer (tail portion of Figure 1g left), consistent with its fast dissociation from a binary complex with DNA.^47^

The coincidence of condensate spreading and strand reannealing during retraction and the reformation of foci in the subsequent overstretch were also clearly captured by kymographs (Figures 1h and S2b). Each focus migrated in one direction, toward the adjacent ssDNA section. With the higher time resolution of kymographs, one could even discern the trace behind the migrating focus before complete spreading. It was as if the focus moved together with the advancing ss-dsDNA junction in spreading the condensate. After one retract-stretch cycle in buffer, dipping the λ-DNA tether in the HMGB1 channel revived foci (Figure S2b far right). In an alternative protocol, we soaked a relaxed λ-DNA tether (extension ∼7 µm) in the HMGB1 channel and moved it outside (Figure S2c). Initially, a uniform layer of HMGB1 was spread over the DNA surface. Upon stretching the DNA tether, the spread converted into distinct foci, confirming the reversibility of the focus-spread transition.

To unequivocally determine the localization of HMGB1 foci, we stained λ-DNA with YOYO-3, a dsDNA intercalating dye (Figure S2d). On overstretched λ-DNA, gaps between YOYO-3-stained sections represented unpeeled ssDNA sections. HMGB1 foci always occurred at ss-dsDNA junctions. Interestingly, there was an apparent correlation between the size of focus and the length of the corresponding peeled ssDNA section, suggesting that each peeled-off ssDNA track was entirely drawn into a focus. The two observations, ready spread of HMGB1-mediated condensates and coincidence of condensate spreading and strand reannealing, together suggest that co-condensation at ss-dsDNA junctions promotes strand reannealing. Indeed, overstretched λ-DNA often showed residual strand separation in buffer (Figures 1i) but predominantly showed complete reannealing in the presence of HMGB1 (Figures 1j); 5 µM HMGB1 increased the probability of complete reannealing from 0.36 to 0.85 (Figures 1k). In contrast, reannealing of overstretched λ-DNA was always incomplete in the presence of protamine (Figure 1d top and additional data in refs ^16, 30^).

In short, while both HMGB1 and protamine produced foci at ss-dsDNA junctions, their actions exhibited stark contrasts. First off, HMGB1 foci were observed only in overstretch whereas protamine foci persisted to low stretch. Related to that, force–extension curves of protamine-bound λ-DNA revealed hyperstable tangles that withstood forces > 60 pN (Figure 1d bottom), but HMGB1 did not condense dsDNA and only increased its flexibility (Figures 1f and S1). Moreover, HMGB1 foci promoted DNA strand separation but protamine foci hindered it. Lastly, unique to HMGB1, foci spread at low stretch, which is a wetting transition on the dsDNA surface. This transition was accompanied by strand reannealing, such that pre-transition condensates were dominated by ssDNA but post-transition spread solely involved dsDNA. As noted above, protamine-mediated condensates initiated on overstretched DNA also underwent a transition from an ssDNA-dominated state to a dsDNA-dominated state. However, unlike HMGB1, protamine-mediated condensates in the dsDNA-dominated state did not wet the DNA surface; on the contrary, they drew in more and more dsDNA (Figure 1a-c). As we concluded previously, protamine interacts more strongly with dsDNA than with ssDNA.^30^ In contrast, as detailed below, HMGB1 interacts more strongly with ssDNA than with dsDNA in co-condensation. The weaker interaction with dsDNA explains the spread of HMGB1-mediated condensates.

### Counteraction of HMGB1 converted protamine-mediated tangles into bridges

We now present results from retracting and stretching λ-DNA in the simultaneous presence of HMGB1 and protamine. We followed a general protocol (Figure S3a) in which an overstretched λ-DNA tether was brought into the HMGB1 + protamine channel for soaking or retract-stretch cycles and then brought outside for one more retract-stretch cycle or confocal scanning. Based on the observation that both HMGB1-only and protamine-only foci localized at ss-dsDNA junctions, we anticipated that HMGB1 foci and protamine foci would colocalize when both proteins were present. 2D scans indeed confirmed this colocalization (Figure 2a).

**Figure 2.**
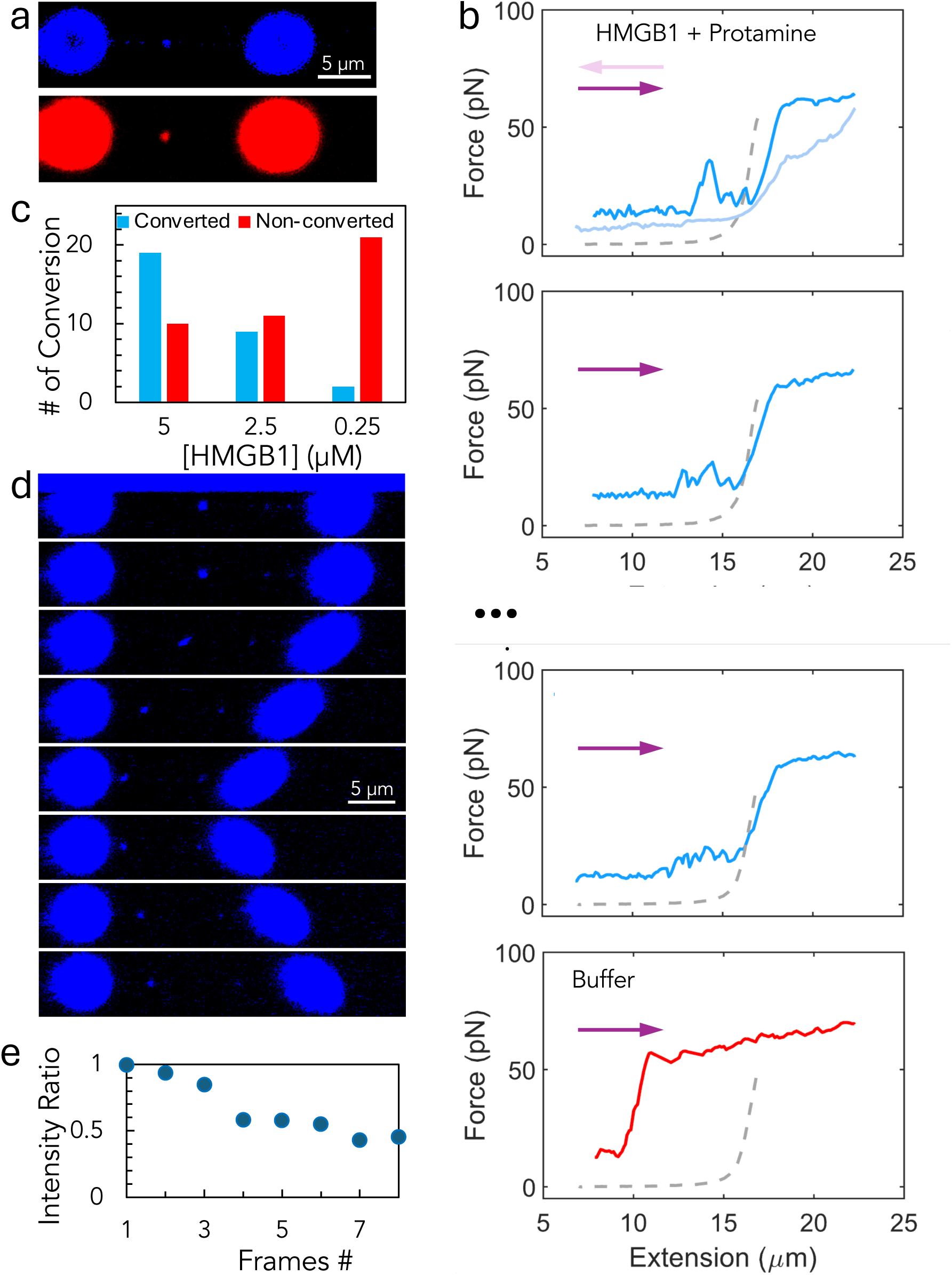
HMGB1’s colocalization with and counteraction to protamine. (**a**) 2D scans showing colocalization of PM-Cy5 (red) and GFP-HMGB1 (blue) in a focus formed on λ-DNA. An overstretch λ-DNA tether was soaked in the HMGB1 + protamine channel and then retracted to 15 µm in the buffer just outside for scanning (see Figure S3a). Protein concentrations were (in µM): HMGB1, 8; GFP-HMGB1, 0.8; protamine 10; protamine-Cy5, 1.25. (**b**) Force-extension curves of λ-DNA during consecutive retract-stretch cycles in a channel containing 5 µM HMGB1 and 2.5 µM protamine. Two cycles are not shown (indicated by ellipses). The last cycle (curve color changed from blue to red) was carried out in the buffer just outside the channel (see Figure S3a). (**c**) A bar graph showing the number of λ-DNA tethers for which tangles were or were not converted into bridges in the HMGB1 + protamine channel, at the indicated HMGB1 concentrations. Protamine was fixed at 2.5 µM. Non-conversion meant that the first three stretching curves all showed tangles whereas conversion meant at least one of the three curves showed bridges. (**d**) 2D scans of a λ-DNA tether undergoing retraction and stretching immediately after moving outside the HMGB1 + protamine channel (8 µM HMGB1; 0.8 µM HMGB1; and 2.5 µM protamine). (**e**) Fluorescence intensities in the major focus (one on the left) of (**d**), normalized by the value in the first frame.

Given their contrasting effects delineated above, we further anticipated that HMGB1 would counteract the effects of protamine. The signature action of protamine is to produce tangles initiated at ss-dsDNA junctions. When HMGB1 was also present, tangles no longer formed, as the rise to strand separation occurred at the same or a longer extension when compared to the counterpart of naked DNA (Figures 2b and S3b, c, curves in blue color). Instead of tangles, bridges, characterized by a sawtooth pattern with rupture forces at ∼25 pN, along with bends at ∼10 pN were observed. The tangle-to-bridge conversion rate by HMGB1 was dose-dependent (Figure S4), decreasing from 66% at 5 µM to 9% at 0.25 µM (with protamine fixed at 2.5 at a fixed; Figure 2c). Out of the 19 tethers on which tangle-to-bridge conversion occurred at 5 µM HMGB1, 10 (or 52%) were accompanied by complete strand reannealing. The addition of ssDNA could convert protamine-mediated tangles into bridges.^30^ Here we found that HMGB1 could also produce this conversion without any help from ssDNA. When the λ-DNA tethers were finally brought outside the HMGB1 + protamine channel, tangles reappeared (Figures 2b and S3b, c, curves in red color). Previously we have shown that protamine remained bound when DNA tethers were transferred to buffer.^16^ The reappearance can be attributed to the dissociation of HMGB1 into buffer (see fluorescence attenuation in the tail portion of Figure 1g left). 2D scans of a tether brought just outside the HMGB1 + protamine channel confirmed fluorescence attenuation and hence HMGB1 dissociation (Figure 2d, e). These scans further showed that, instead of spreading, HMGB1 remained inside foci at low stretch. As these scans were taken outside the HMGB1 + protamine channel and thus could be complicated by HMGB1 dissociation, we also took scans when a λ-DNA tether was inside the channel, confirming HMGB1 residence in foci (Figure S3d). So the conversion of solid-like tangles to liquid-like bridges by HMGB1 is due to its action inside condensates, mediated by, e.g., interaction with protamine.

We also experimented with protocols in which HMGB1 and protamine were placed in different channels. The force-extension curve of an intact-DNA tether showed bends in the protamine channel; bends then persisted even after the tether was moved into the HMGB1 channel (Figure S5a), reflecting the dominance of protamine that remained bound to the DNA. When the order of going to the two channels was reversed, the force-extension curves showed the expected increase in dsDNA flexibility in the HMGB1 channel but then bends in the protamine channel (Figure S5b). 2D scans revealed that an initial layer of HMGB1 on the DNA disappeared when the tether was moved to the protamine channel (Figure S5c), due to HMGB1 dissociation. Similarly, an HMGB1 focus formed on an overstretched DNA tether lost fluorescence after moving to the protamine channel (Figure S5d). Correspondingly, force-extension curves showed persistence of tangles (Figure S5e). These results demonstrate that the counteraction of HMGB1 needed an HMGB1 pool to maintain its presence in protamine-mediated condensates.

### Counteraction of HMGB1 required its acidic tail

We wondered whether the acidic tail of HMGB1 was required for its counteraction against protamine, and addressed this question using the deletion construct, HMGB1-ΔC. We first designed a competition experiment to demonstrate that HMGB1-ΔC could displace HMGB1 from foci. On overstretched λ-DNA tethers, 0.25 μM GFP-HMGB1 always formed foci (Figure 3a top left), as expected from the results already reported in Figure 1e, h. When 0.25 μM HMGB1-ΔC was also present, GFP-HMGB1 foci were observable only in some scans of a tether (Figure 3a bottom left). Similarly, on intact λ-DNA, GFP-HMGB1 always formed a layer (Figure 3a top right), but a layer was no longer observable in some scans in the presence of 0.25 μM HMGB1-ΔC (Figure 3a bottom right). The probability of observing foci decreased with increasing HMGB1-ΔC concentration (Figure 3b top), confirming that the apparent disappearance of foci was due to the displacement of GFP-HMGB1 by HMGB1-ΔC (Figure 3b bottom). The HMGB1-ΔC concentration required to reduce the probability of observing foci to 0.5 was 0.21 μM, a little lower than the GFP-HMGB1 used for focus formation. HMGB1-ΔC was thus effective in partitioning into foci at ss-dsDNA junctions.

**Figure 3.**
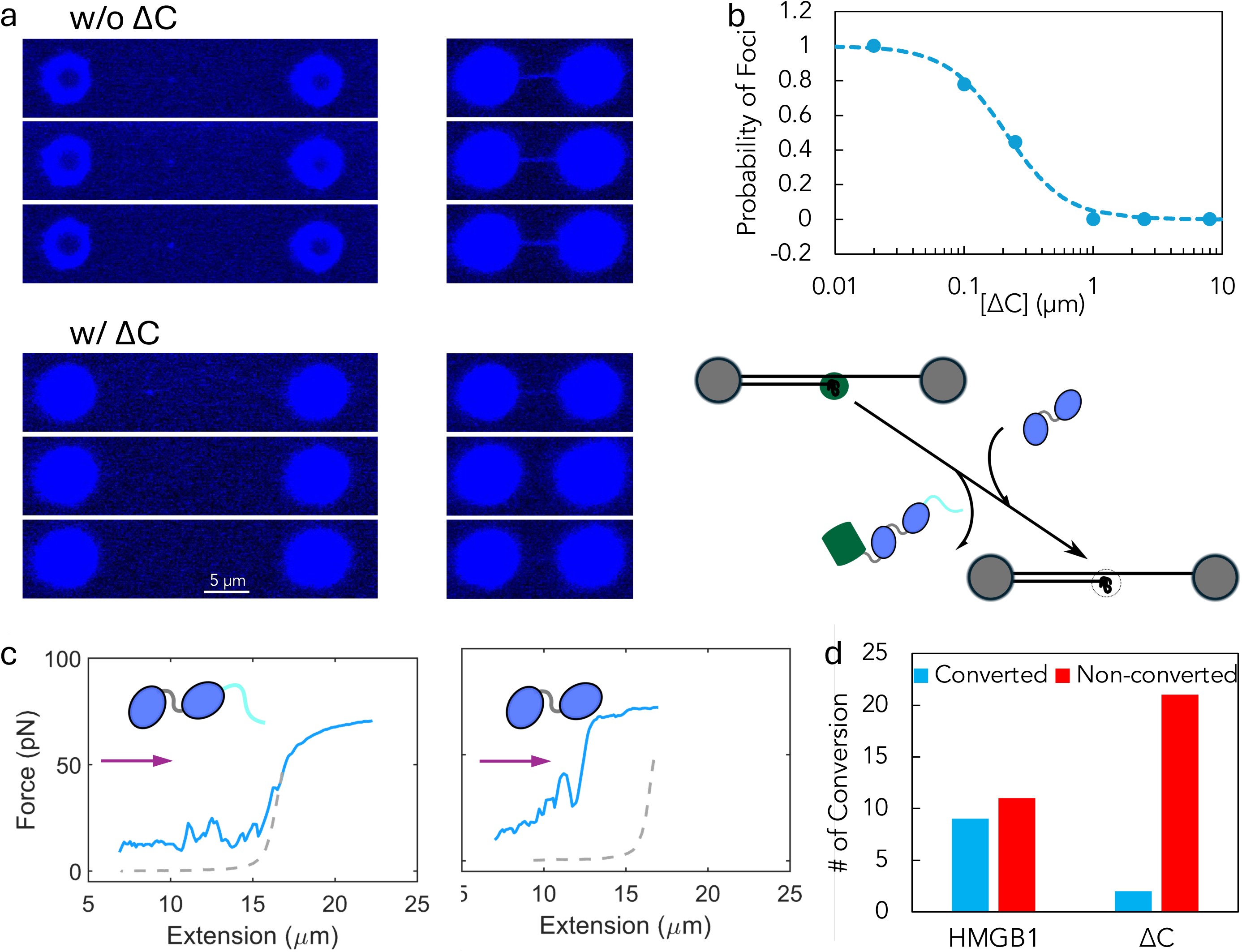
Inability of HMGB1-ΔC in converting protamine-mediated tangles to bridges. (**a**) Top: 2D scans of overstretched (extension ∼23 μm; left panels) and intact (extension ∼ 7 μm; right panels) λ-DNA inside a channel containing 0.25 μM GFP-HMGB1. Bottom: corresponding scans when the channel also contained 0.25 μM HMGB1-ΔC. A tether was scanned three times after moving into the channel under each condition; foci (left panels) or a layer (right panel) was visible in all three scans without HMGB1-ΔC but only visible in the first scan with HMGB1-ΔC. (**b**) The probability of observing HMGB1 foci in a 2D scan when HMGB1-ΔC was present together with 0.25 μM GFP-HMGB1. Probabilities were averaged over three consecutive scans of each tether and then over three tethers at each HMGB1-ΔC concentration. Bottom: schematic illustrating the exchange between GFP-HMGB1 and HMGB1-ΔC in foci. (**c**) Force-extension curves showing conversion of tangles to bridges by 2.5 μM (left) but failure to do so by 2.5 μM HMGB1-ΔC (right) when protamine was present at 2.5 µM. (**d**) A bar graph showing the number of λ-DNA tethers for which tangles were or were not converted into bridges in the HMGB1(-ΔC) + protamine channel (2.5 µM protamine along with 2.5 μM HMGB1 or HMGB1-ΔC). The HMGB1 results are the same as in Figure 2c.

We performed a similar competition experiment using HMGB1 (Figure S6). GFP-HMGB1 foci on overstretched λ-DNA faded with increasing HMGB1concentrations, but remained visible at 10 μM and only disappeared at 25 μM. The last two values bracket the HMGB1 concentration at which the probability of observing foci was 0.5. This HMGB1 concentration was much higher than the corresponding value, 0.21 μM, for HMGB1-ΔC, reflecting weakened interactions with DNA by the acidic tail.

Next we addressed whether HMGB1-ΔC was competent in converting protamine-mediated tangles into bridges by collecting force-extensions of initially overstretched λ-DNA in the presence of both protamine and HMGB1-ΔC (Figure 3c). The tangle-to-bridge conversion rate was reduced from 45% for 2.5 µM HMGB1 to only 9% for 2.5 µM HMGB1-ΔC (Figure 3d). Interestingly, 2.5 µM GFP-HMGB1 changed the tangle-to-bridge conversion rate in the opposite direction, raising it to 88% (Figure S7). GFP tags affected the condensation of many proteins, with the level of the tags’ negative charges playing a major role.^52^ To avoid GFP-related artifacts, we limited to 10% GFP-HMGB1 doping for imaging and collected most force-extension curves without GFP-HMGB1. We also confirmed focus formation on overstretched λ-DNA using HMGB1 labeled with a small fluorescent molecule Cy3 (Figure S8).

With the foregoing results with HMGB1-ΔC and GFP-HMGB1, we can now be more definitive in the mechanism underlying the conversion of protamine-mediated solid-like tangles to liquid-like bridges. HMGB1 molecules, via their acidic tails, likely peel some protamine molecules away from their very strong interactions with dsDNA and form weaker interactions with them instead, leading to condensate liquidity. The acidic tail of HMGB1 is essential for interactions with protamine in co-condensation (see below); its deletion impairs HMGB1’s ability to compete with dsDNA for protamine, hence the substantial reduction in tangle-to-bridge conversion rate (Figure 3d). On the other hand, the net negative charge (–6) of the GFP tag enhances the electrostatic attraction of the acidic tail to protamine and therefore the competitiveness of HMGB1 against dsDNA for protamine, explaining the elevated tangle-to-bridge conversion rate of GFP-HMGB1 (Figure S7).

### HMGB1 possessed a higher co-condensation propensity with ssDNA than with dsDNA

We performed a series of bulk experiments to cross-validate the single-molecule results and provide additional mechanistic insight. First, we used brightfield microscopy to check the co-condensation propensities of HMGB1 with 25-bp dsDNA or 32-nt ssDNA. HMGB1 and dsDNA did not co-condense even at very high concentrations (100 µM each), but HMGB1 and ssDNA readily formed liquid droplets (Figure 4a top). This result aligns precisely with the single-molecule observations that HMGB1 formed foci with peeled-off ssDNA tracks on overstretched λ-DNA but did not condense dsDNA at low stretch, forming a layer instead (Figure 1e, f). We can thus now conclude that HMGB1 interacts more strongly with ssDNA than with dsDNA in co-condensation. Histone H1 similarly exhibited a higher co-condensation propensity with ssDNA than with dsDNA.^50^ In contrast, protamine co-condensed into amorphous aggregates with dsDNA but into liquid droplets with ssDNA, and induced solid-like tangles on single dsDNA but liquid-like bridges on single ssDNA.^16, 30^ Both observations point to stronger interactions of protamine with dsDNA than with ssDNA, which can be attributed to prevalent arginine-nucleobase hydrogen bonding according to all-atom molecular dynamics simulations.^16^

**Figure 4.**
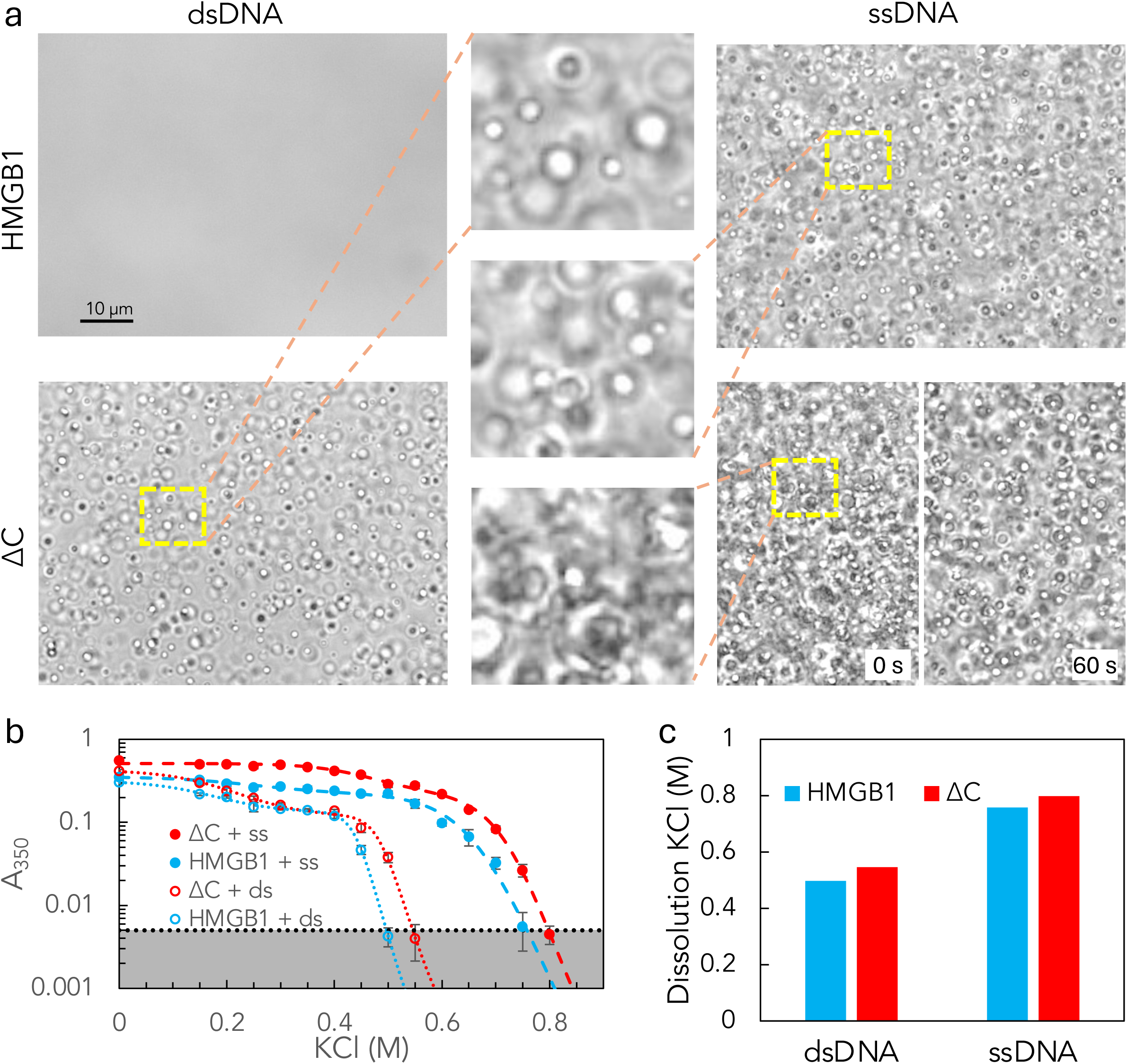
Co-condensation propensities of HMGB1 and HMGB1-ΔC with DNA. (**a**) Brightfield images of HMGB1 or HMGB1-ΔC mixed with 25-bp dsDNA (left) and 32-nt ssDNA (right). The middle column displays zoomed views of regions in a yellow box. Images were taken immediately after loading onto slides (0 s). For the ΔC–ssDNA mixture, the image at 0 s shows very crowded droplets; an image at 60 s is also displayed, showing droplets at a slightly reduced density due to sedimentation. All components were at 100 µM, mixed in 10 mM imidazole buffer (pH 7). (**b**) Effects of salt on droplet population, as measured by absorbance at 350 nm (A_350_). Error bars display standard deviations among four replicates. Proteins and 25-bp dsDNA were each at 20 µM; 32-nt ssDNA was at 31.25 µM (for charge matching with dsDNA); all were dissolved 10 mM imidazole buffer (pH 7) with 10% PEG. Curves are a fit to equation [5]. The shaded region indicates background levels. (**c**) The lowest KCl concentrations at which droplets were dissolved.

Unlike HMGB1, HMGB1-ΔC co-condensed into liquid droplets with both dsDNA and ssDNA (Figure 4a bottom). With ssDNA, the number of droplets formed by HMGB1-ΔC was much higher than by HMGB1. So the interactions of HMGB1 with both forms of DNA were strengthened upon deleting the acidic tail, which could interfere with DNA binding by occupying the HMG boxes^45–47^ and by electrostatic repulsion. Both the single-molecule (Figures 3b and S6a) and bulk experiments thus captured the expected effects of the acidic tail. By comparing the number of droplets, we can also conclude that, like HMGB1, HMGB1-ΔC interacts more strongly with ssDNA than with dsDNA.

We used salt dissolution of droplets to confirm the difference between HMGB1 and HMGB1-ΔC in co-condensation. For this experiment, we formed droplets of all four binary mixtures by including 10% PEG, and added increasing levels of KCl to dissolve them. The amounts of droplets were quantified by absorbance at 350 nm (Figure 4b). For both dsDNA and ssDNA, the amounts of droplets decayed more slowly to 0 for HMGB1-ΔC than for HMGB1. The dissolution KCl concentrations were 0.50 and 0.55 M, respectively, for HMGB1 and HMGB1-ΔC mixtures with dsDNA, and 0.76 and 0.80 M, respectively, for the corresponding mixtures with ssDNA (Figure 4c).

### HMGB1 but not HMGB1-ΔC converted protamine-dsDNA aggregates into liquid droplets

Given the single-molecule observation that HMGB1 converted protamine-mediated tangles into bridges, we tested the ability of HMGB1 to convert protamine–dsDNA aggregates into droplets in a bulk setting. A mixture of 10 µM protamine and 20 µM 25-bp DNA formed small aggregates (Figure 5a). Upon adding 20 µM HMGB1 (along with 10% PEG), aggregates were converted into droplets with a homogeneous distribution of both proteins (Figure 5b). In contrast, repeating the experiment with HMGB1 replaced by HMGB1-ΔC failed to produce conversion. In fact, aggregates grew more numerous and bigger with the addition of HMGB1-ΔC (Figure 5c).

**Figure 5.**
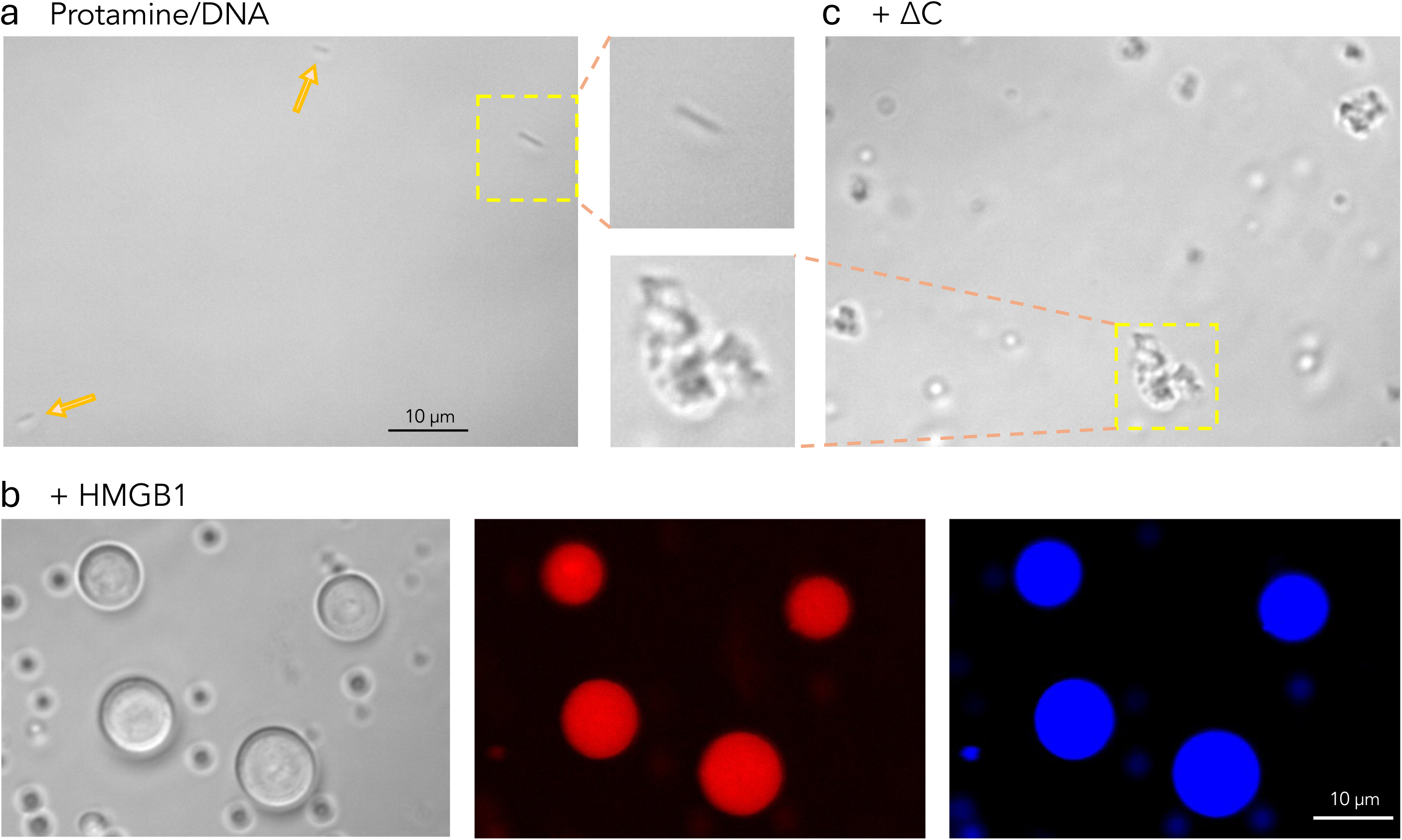
Conversion of protamine–dsDNA aggregates into droplets by HMGB1. (**a**) Brightfield image of a protamine–dsDNA mixture, showing sparse, small aggregates (indicated by arrows and box). Concentrations were: protamine, 9 µM; protamine-Cy5, 1 µM; 25-bp dsDNA, 20 µM; mixed in 10 mM imidazole buffer (pH 7). (**b**) Brightfield and confocal images of a ternary mixture, with 19 μM HMGB1 and 1 μM GFP-HMGB1 (plus 10% PEG) added to preformed protamine–dsDNA aggregates described in (**a**), showing droplets with a homogenous distribution of both protein components. Four droplets were grown into large sizes by using an optical trap to scavenge small droplets. (**c**) Brightfield image of a ternary mixture, with 20 μM HMGB1-ΔC (plus 10% PEG) added to preformed protamine–dsDNA aggregates, showing aggregates with increased number and size relative to (**a**).

These observations support our mechanistic interpretation of single-molecule results on tangle-to-bridge conversion. When forming liquid droplets in the ternary mixture, HMGB1 indeed appears to peel a portion of protamine molecules away from dsDNA. The need for 10% PEG in the bulk experiment could be due to an effective dilution when the dimensionality of space is increased from one (on the dsDNA in the single-molecule setting) to three (in solution); HMGB1 required PEG for homotypic condensation in solution (ref ^49^; Figure S9a) but formed a layer on single dsDNA chains without PEG. When the acidic tail is deleted, the ability to peel away protamine is impaired. This assertion is validated by the observation of droplet formation by an HMGB1–protamine mixture and lack of co-condensation when HMGB1 was replaced by HMGB1-ΔC (Figure S9b). Instead, the HMGB1-ΔC construct reinforces the strong interaction of protamine with dsDNA, leading to the observed increase in number and size of aggregates in Figure 5c.

### Protamine slowed down the dynamics of HMGB1–dsDNA droplets

HMGB1 formed a layer on dsDNA without condensing it (Figure 1e, f); when protamine was added, dsDNA was condensed into bridges (Figure 2b). Intermolecular interactions formed by protamine, particularly strong with dsDNA and somewhat weaker with HMGB1, are expected to have a major impact on condensate dynamics. As HMGB1 and dsDNA could co-condense in the presence of 10% PEG (Figure 4b), we compared the internal dynamics of HMGB1–dsDNA (HMG/DNA for short) and HMGB1–dsDNA–protamine droplets. Ternary droplets were prepared in two ways: first forming HMG/DNA droplets with PEG and then adding protamine (denoted as HMG/DNA + PM), or first forming protamine–dsDNA aggregates and then adding HMGB1 along with PEG (denoted as PM/DNA + HMG).

We first used FRAP to probe internal dynamics (Figure 6a). Fluorescence recovery was near complete (∼95%) in both binary droplets and the two preparations of ternary droplets (Figure 6a, b). Recovery was slower in ternary droplets (Figure 6c). From fitting to an exponential function, the recovery times were 9.3 ± 1.3 s in binary droplets and 12.4 ± 1.5 s and 11.6 ± 0.3 s in the two preparations of ternary droplets (Figure 6d). The addition of protamine thus slowed down fluorescence recovery very significantly (*p* value < 0.01).

**Figure 6.**
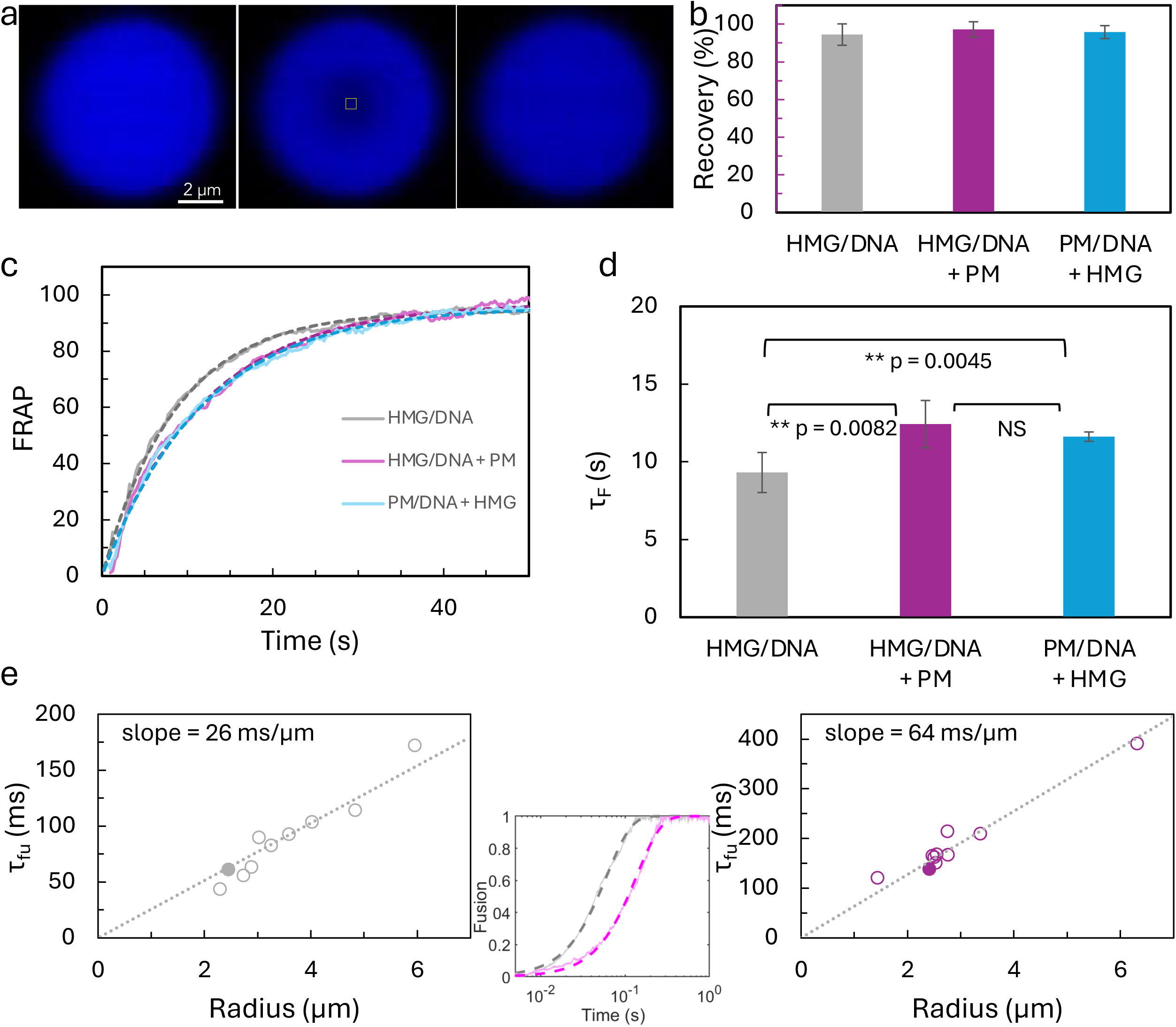
FRAP in and OT-directed fusion of droplets formed by binary and ternary mixtures of HMGB1, dsDNA, and protamine. (**a**) Confocal images showing an HMGB1–dsDNA droplet before bleach, immediately after bleach (0 s), and 60 s post-bleach. A yellow box indicates the bleach region. Droplets were formed by mixing 19 μM HMGB1, 1 μM GFP-HMGB1, and 20 µM 25-bp dsDNA in 10 mM imidazole buffer (pH 7) plus 10% PEG. (**b**) Percentages of fluorescence recovery in three types of droplets: HMG/dsDNA, binary mixture as described in (**a**); HMG/dsDNA + PM, 10 µM protamine added to binary droplets; PM/dsDNA + HMG, ternary mixture as described in Figure 5b. Fluorescence recovery was calculated as the ratio of intensities in the bleached region and a background region at 60 and 200 s, respectively, post-bleach for binary and ternary droplets. Error bars display standard deviations among five replicates. (**c**) Representative fluorescence recovery traces. Solid curves are fits to equation [3]. (**d**) Fluorescence recovery times. Error bars display standard deviations among five replicates. (**e**) Left: fusion time of HMG/dsDNA droplets as a function of initial droplet radius. Each circle displays the fusion time of a pair of droplets; the line displays proportionality between fusion time and droplet radius; right: corresponding results for HMG/dsDNA + PM droplets. Middle: fusion progress curves of a pair of binary droplets (gray) and a pair of ternary droplets (magenta). Dashed curves are fits to equation [4]. The corresponding 1_fu_ values are shown as solid circles in the panels at left and right.

OT-directed fusion revealed a similar difference in dynamics between HMG/DNA and HMG/DNA + PM droplets (Figure S10). The inverse fusion speed was 26 ms/µm for binary droplets and increased to 64 ms/µm for ternary droplets (Figure 6e). A correlation between fluorescence recovery time and inverse fusion speed was observed previously.^53^

## Discussion

We have shown that, like protamine, HMGB1 forms foci at ss-dsDNA junctions on overstretched single DNA. However, the actions of these two proteins otherwise exhibit multiple contrasts, including dsDNA condensation and maintenance of foci versus absence of dsDNA condensation and focus spreading at low stretch, resistance versus promotion of strand reannealing, and formation of hyperstable tangles versus a mere increase in DNA flexibility (Figure 7). When both proteins are present, condensates initiated at ss-dsDA junctions comprise liquid-like bridges and still promote strand reannealing. The acidic tail of HMGB1 is indispensable for condensate liquidity. The bulk experiments have provided additional mechanistic insights, including the slowdown of condensate dynamics by strong protamine-dsDNA interactions.

**Figure 7.**
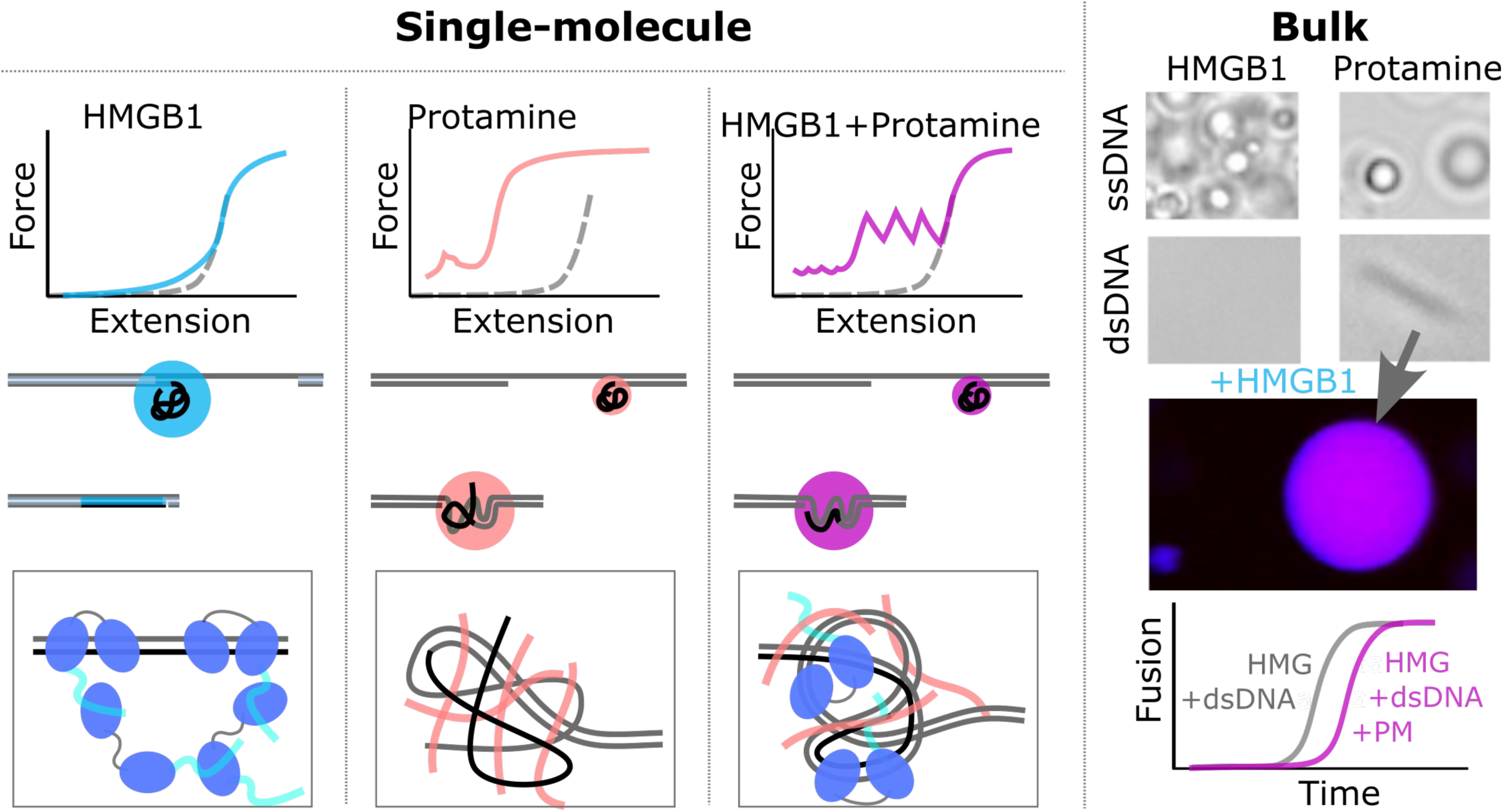
A summary of the contrasting actions of HMGB1 and protamine and the effects of their simultaneous presence. The bottom middle rows in the single-molecule portion display overstretched or relaxed λ-DNA, respectively, where peeled-off ssDNA tracks are in dark while ssDNA and dsDNA sections under tension are in gray.

In forming a layer on the surface of relaxed dsDNA, the HMG boxes of HMGB1 molecules would have to bind sites along the DNA (Figure 7, last row in the single-molecule portion). The freed acidic tails would then bind HMG boxes of additional HMGB1 molecules, and this process may propagate multiple rounds. With overstretched λ-DNA, a thin coating of HMGB1 instead of a layer was observed on dsDNA sections (Figures 1e and S8), because HMGB1 likely has a higher affinity for relaxed dsDNA than for stretched dsDNA. The acidic tail can of course bind an HMG box on the same molecule, leading to its dissociation from the surface layer. This intramolecular binding explains the ready dissociation of HMGB1 from dsDNA when a tether was moved outside the HMGB1 channel. In contrast, protamine can crosslink both ssDNA and dsDNA; condensates are liquid-like when dominated by ssDNA but solid-like when dominated by dsDNA.^30^ When both proteins are present in a ternary condensate with dsDNA, acidic tails of HMGB1 molecules peel protamine molecules away from dsDNA, leading to condensate liquidity.

While monomeric GFP (mGFP) is often thought of as an inert reporter,^54^ mGFP tags have recently been found to affect the condensation of many proteins, with the tags’ net negative charges playing a major role.^52^ Here we have shown that our mEGFP tag affects the material property of a ternary condensate composed of HMGB1, protamine, and dsDNA. Specifically, the mEGFP tag, via its net negative charge (–6), reinforces the acidic tail of HMGB1 in peeling protamine away from dsDNA, thereby making the condensate more liquid-like, as indicated by an elevated tangle-to-bridge conversion rate. The opposite effects of the mEGFP tag and the acidic-tail deletion on tangle-to-bridge conversion were very helpful in developing a mechanistic understanding of HMGB1’s counteraction against protamine.

In nucleosome-based chromatin, HMGB1 counters linker histone H1 and increases the accessibility of nucleosomal DNA, for transcription and DNA repair.^35, 36^ Binding, FRAP-based live-cell competition, and biochemical studies suggest that the acidic tail of HMGB1 binds with the basic tail of H1, thereby loosening the latter’s interaction with DNA.^33, 34, 37^ This mechanism is similar to one we developed here for the counteraction of protamine against HMGB1 (Figure 7, last row in the single-molecule portion). There is much in common between histone H1 and protamine. H1 has a highly basic tail, similar to the arginine-rich composition of protamine. In a previous single-molecule study, both proteins were found to condense ssDNA into liquid-like bridges.^30^ Both proteins have much higher affinities for DNA (or nucleosomes) than HMGB1 (sub-nm^18, 55^ versus 100 nM^37^), although this measure of binary interactions may not be strictly relevant for polyvalent interactions in molecular networks such as in condensates.^56^ The interplay between DNA-binding proteins may be a general mechanism for fine-tuning the dynamics of DNA condensates.

HMGB1 formed foci at ss-dsDNA junctions on overstretched λ-DNA, which then spread upon retraction, accompanied by strand reannealing. Spreading is a wetting transition on the dsDNA surface, distinct from a prewetting transition observed with the transcription factor Klf4.^57^ Prewetting transitions are between a thin coating and a thick layer, whereas wetting transitions are between a focus, representing a partial wetting state, and a layer on the DNA surface, representing a complete wetting state. The wetting transition of HMGB1 occurred at the same time as the strands were reannealed, so that the pre-transition focus was dominated by ssDNA but the post-transition spread solely involved dsDNA. Kymographs (Figure S2b) suggest that a focus migrates in the direction of the neighboring unpeeled ssDNA section, leaving a layer of HMGB1 behind. It thus appears that reannealing proceeds through the zipping of the two strands at the ss-dsDNA junction, with the focus moving with the advancing junction and leaving behind a layer on the newly reannealed section. Focus formation and strand reannealing have a symbiotic relationship: a focus holds the peeled-off ssDNA track in close proximity to the ss-dsDNA junction for zipping, while reannealing reduces the amount of ssDNA in the focus, thereby weakening it for spreading. We speculate that the foregoing properties of HMGB1 facilitate its function in DNA repair:^44^ focus localization at ss-dsDNA junctions may provide a mechanism for the recruitment of HMGB1 to damage sites, and promotion of strand reannealing may ease the completion of the repair process.

In spermiogenesis of mammals and flies, DNA supercoiling builds up as histones are removed; topoisomerase IIβ creates double-strand breaks to relieve supercoiling.^7–9^ At this stage, transition proteins and other chromatin-associated factors including HMGB1 and HMGB2 are transiently expressed as protamines also start to appear.^10–13^ We hypothesize that transiently expressed proteins counter protamines to keep condensed DNA in a liquid state, in order to facilitate the recruitment of DNA repair machinery to restore duplex structure. In mammals, transition proteins are Arg/Lys-rich whereas protamines are exclusively Arg-rich. Arg sidechains can uniquely form hyperstable wedges with nucleobases;^16^ Arg-to-Lys substitutions in protamines significantly reduced DNA binding affinity.^18^ Consequently, whereas co-condensation of dsDNA with protamine produced solid-like aggregates,^16, 26^ co-condensation with transition protein TNP1 produced liquid droplets.^58^ Once transition proteins are degraded and DNA duplex structure is restored, extreme chromatin compaction by protamines leads to global silencing of transcription. In flies, spermiogenesis is mediated by SNBPs, which contain both one or two HMG boxes and Arg/Lys-rich motifs, and can be separated into transition protein-like and protamine-like (Table S1). The present work suggests that competition between HMG boxes and Arg/Lys-rich motifs for DNA binding allows SNBPs to tune the dynamics of condensed chromatin; SNBPs eliciting a higher level of dynamics thus act as transition proteins while those eliciting a lower level of dynamics act as protamines. In support of this suggestion, transition protein-like SNBPs contain a low number (3 to 10) of Arg and Lys residues in their Arg/Lys-rich motifs, whereas protamine-like SNBPs contain a high number (13 to 23) of Arg and Lys residues, thereby enabling stronger DNA binding and slower condensate dynamics. In embryonic development, a reverse, histone-to-protamine transition requires DNA decondensation, which is triggered by protamine phosphorylation.^26^ HMGB1 is expressed in preimplantation embryos;^59, 60^ whether it plays a role in DNA decondensation there remains to be studied.

## Materials and Methods

### Materials

Protamine sulfate (catalog # P4020) was from Sigma Aldrich. Salt was removed by dialysis ^61^ before use. SYTOX Orange (catalog # 00012), streptavidin-coated polystyrene beads (4.35 µm diameter; catalog # 00034-6), and biotinylated 48.5 kb λ-DNA (catalog # 00001) were from LUMICKS. YOYO-3 (catalog # Y3606) was from Thermo Fisher Scientific. 25-bp dsDNA and 32-nt ssDNA were custom-ordered from IDT. The sequence of the first strand in the 25-bp dsDNA was 5’-ACGTTATGGC AGTCGTTAAA TTGAG-3’ For the 32-nt ssDNA, the sequence was 5’-GAAACGCTCA GGGAAGAAGA AGATCAGCAA AG-3’ Imidazole buffer (10 mM pH 7) was prepared by dissolving imidazole (catalog # 396745000, Thermo Scientific) in Milli Q water. PEG 6k (catalog # 81255) was from Sigma Aldrich. Unless otherwise stated, 150 mM KCl was added to make the working buffer. All solutions were filtered using 0.22-μm filters (catalog # SLGPR33RS, Millipore Sigma). All experiments were done at room temperature.

### Protein expression and purification

His-tagged HMGB1 and HMGB1-ΔC plasmids were gifts from Dr. Geeta Narlikar at UC San Francisco. Expression and purification followed a published protocol^37^ with minor modifications. Briefly, after transformation of the plasmids, Rosetta (DE3) competent cells were grown in LB media supplemented with 50 µg/mL carbenicillin and 25 µg/mL chloramphenicol at 37 °C. Upon reaching an OD of 0.6, expression was induced by IPTG (1 mM) at 37 °C for 4 h. Cells were pelleted by spinning for 30 minutes at 5,000g at 4 °C. The cell pellet was resuspended in lysis buffer (20 mM HEPES-KOH pH 7.5, 500 mM KCl, 10% glycerol, 2 mM BME, 2 mM PMSF, 7.5 mM imidazole, 2 µg/mL pepstatin A, 6 µg/mL leupeptin, and 4 µg/mL aprotinin), and lysed using a homogenizer (EmulsiFlex-C5, Avestin) at 15,000 psi, repeated four times, followed by spinning at 40,000g for 30 minutes at 4 °C. The supernatant was filtered using 0.25 µm filters and loaded onto a HiTrap TALON crude column (Cytiva, catalog # 28953767), which was equilibrated with lysis buffer. The column was washed with lysis buffer without protease inhibitors, and eluted using elution buffer (20 mM HEPES-KOH pH 7.5, 150 mM KCl, 2 mM BME, 400 mM imidazole). After adding TEV protease (150 μg of TEV/liter of bacterial culture) to cleave the His-tag, the eluate was dialyzed overnight at 4 °C in dialysis buffer (20 mM HEPES-KOH pH 7.5, 150 mM NaCl, 2 mM dithiothreitol). After dialysis, the protein was loaded to a HiTrap Q HP (for HMGB1) or SP HP (for HMGB1-ΔC) column (Cytiva, catalog #s 17115401 and 17115201) equilibrated with 10% QB buffer and eluted in a linear gradient (10% to 60%) of QB buffer (20 mM HEPES-KOH pH 7.5, 3 mM dithiothreitol, 1.5 M KCl; percentage was specifically for KCl). Fractions containing the protein were pooled, concentrated using a 3000-Da MWCO Amicon ultra centrifugal filter (Millipore Sigma, catalog # UFC800324), and buffer exchanged (50 mM Tris-HCl, 0.5 mM dithiothreitol) for storage at −80 °C.

The His-tagged mEGFP–HMGB1 plasmid was from Addgene (catalog # 194543^49^) and transformed in Rosetta (DE3) pLysS cells (Sigma-Aldrich). Expression and purification proceeded similarly as just described. Cells were grown in LB media with 100 µg/mL ampicillin and 25 µg/mL chloramphenicol at 37 °C followed by IPTG induction (1 mM) overnight at 25°C. Pellets were resuspended in ice-cold lysis buffer [0.1 % Triton X-100, 5 % glycerol, and protease inhibitor tablets (Roche)], and lysed using a homogeniser at 15,000 psi for 30 min. The lysate was centrifuged at 40,000g for 30 min at 4 °C and the supernatant was loaded onto a HisTrap HP column (Cytiva, catalog # 17524802) equilibrated with buffer A (50 mM Tris-HCl, 150 mM NaCl, 10 mM imidazole, and 1 mM dithiothreitol), and eluted with buffer B (50 mM Tris-HCl, 150 mM NaCl, 250 mM imidazole, and 1 mM dithiothreitol). The eluate was then loaded onto a HiTrap Q HP column equilibrated with 10% buffer QB and eluted in a linear gradient (10% to 65%) of QB buffer. Fractions containing GFP-HMGB1 were pooled and concentrated in a 3000-Da MWCO Amicon ultra centrifugal filter stored at −80°C in storage buffer (50 mM Tris-HCl, 150 mM NaCl, 10 % glycerol, and 3 mM dithiothreitol).

### Labeling of protamine with Cy5 and HMGB1 with Cy3

Protamine (40 µM) was labeled by mixing with a 10-fold molar excess of Cy5-NHS ester (400 µM; Lumiprobe Life Science Solutions, catalog # 13020) in 500 µL of PBS buffer pH 8.5. The reaction mixture was incubated overnight at 4 °C on a tube rotator in the dark, covered by aluminum foil, then eluted on a PD MiniTrap G-25 column (Cytiva, catalog # 28918007), pre-equilibrated with 10 column volumes of PBS pH 7.5. Fractions were collected dropwise in tubes, and dialyzed for the removal of free dye and buffer exchanged in 10 mM imidazole pH 7. The concentration of protamine-Cy5 was determined by Cy5 absorbance (650 nm; at most one dye molecule per protein molecule, at the N-terminus) on a NanoDrop 2000c spectrophotometer (Thermo Scientific).

HMGB1 (100 µM) was labeled with Cy3-NHS ester (1 mM; Lumiprobe Life Science Solutions, catalog # 11020) similarly, except that after collection of labeled fractions from the PD MiniTrap G-25 column, free dye was removed and the sample was concentrated by using a centrifugal concentrator (Sartorius Vivaspin 500; 10-kDa MWCO). Briefly, the centrifugal concentrator was pre-blocked with 5% glycerol in PBS pH 7.5 for 5 min, followed by spinning at 10,000g for 5 min. Fresh PBS pH 7.5 was added to the concentrator, followed by spinning at 10,000g for 5 min to remove residual glycerol. The eluted fractions were concentrated (20 min for each 500 µL sample) in the pre-blocked concentrator and, after adding PBS pH 7.5, concentrated again to the previous volume. The protein concentration was determined by measuring tryptophan absorbance (280 nm).

### Single-molecule force spectroscopy

Force-extension curves were acquired using the dual-trap OT module integrated with microfluidics (Figure S2a) in a LUMICKS C-trap instrument, as described previously.^30^ The trapping laser was set to 20% overall power and a 50:50 split between trap 1 and trap 2, resulting in a stiffness of 200 pN/µm for each trap. Sample components (300 to 600 µL each) were loaded into syringes connected to microfluidics channels. The three main channels were filled with streptavidin-coated beads (0.004% w/v in working buffer), biotinylated λ-DNA (160 pg/µL) in working buffer, and working buffer (10 mM pH 7 imidazole with 150 mM KCl), respectively. The fourth, side channel was filled with proteins (HMGB1, GFP-HMGB1, HMGB1-ΔC, HMGB1-Cy3, protamine, protamine-C5, their combinations with each other or with a DNA-staining dye SYTOX Orange or YOYO-3).

All channels were opened at 1 bar pressure for 2 min to uniformly fill the laminar flow cell with the sample components. In the laminar flow, two beads were trapped from the bead channel and then brought to the DNA channel to form a tethered configuration. After one or more tethers were formed between the two beads, the configuration was moved to the buffer channel to avoid further DNA binding. In the buffer channel, bead 2 was fixed and bead 1 was pulled until a single tether remained between the two beads. A stretching curve was acquired in the buffer channel. The tether, at either an overstretched or retracted extension, was then moved to the protein channel. After a mild flow under 0.2 bar pressure was opened briefly for ∼10 s, retract-stretch cycles were repeated inside or outside the channel until the tether broke. The pulling speed was 1 µm/s. All force-extension curves were acquired with HMGB1 without GFP-HMGB1 except for those in Figures 1f, S5e, and S7a.

### 2D confocal scans and kymographs

Correlative force-fluorescence measurements were performed on a LUMICKS C-Trap instrument. Force measurements were as described in the preceding subsection; fluorescence imaging was done on a confocal module of the instrument. For imaging of HMGB1 on λ-DNA, GFP-HMGB1 premixed with HMGB1 was loaded into a side channel without (Figure S2a) or with protamine channel (Figure S3a) unless otherwise stated. λ-DNA tethers were overstretched (to 23-24 μm) in the buffer channel (location 1), then soaked in the protein channel (location 2), and finally imaged outside the channel (location 3) while retracting to ∼7 μm. In Figure S3d and Figure S5c (top), imaging was performed inside the protein channel. HMGB1 and protamine were in separate channels in Figure S5c, d.

For imaging of λ-DNA with protamine, SYTOX Orange (0.5 µM) was premixed into the protamine (10 µM) channel. Imaging was performed inside the channel while retracting the tether from ∼ 24 μm to ∼ 7 μm (Figure 1a, b). For imaging YOYO-3 on λ-DNA, YOYO-3 (5 µM) was added together with GFP-HMGB1 (2.5 µM) and scanning was performed in overstretched positions (24-21 μm) outside the channel (Figure S2d). For colocalization of HMGB1 with protamine, GFP-HMGB1 (2.5 µM) was mixed in a channel with PM-Cy5 (1.25 µM) and PM (10 µM). Initially overstretched DNA was retracted ∼ 15 μm for colocalization scanning outside the channel (Figure 2a). 2D scans were performed using 488, 532 and 638 nm lasers for GFP-HMGB1, SYTOX Orange, and YOYO-3, respectively, with 20% laser power, 0.1 ms pixel dwell time, and 0.1 μm pixel resolution. Scan with HMGB1-Cy3 was performed using the 532 nm laser with similar laser settings. Only one laser was switched on at a time to avoid any interference. Kymographs using GFP-HMGB1 and SYTOX Orange were acquired using 488 and 532 nm lasers, respectively, with 20% laser power. Images were processed using ImageJ.

### Analysis of fluorescence intensity in foci on single λ-DNA

In each retracting frame illustrated by Figure 1b, the fluorescence intensity of SYTOX Orange in a circular region of interest covering a single focus was measured using ImageJ. The corresponding extension was measured as the distance between the edges of the two beads. Intensities were normalized by the maximum intensity on λ-DNA in the first scan before retraction. The data in Figure 1c displaying the dependence of normalized intensity (*I*) on extension (*x*) were fit to

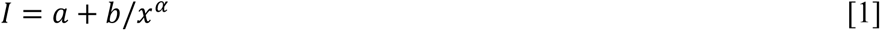

Similarly, HMGB1-GFP intensities in foci illustrated in Figure 1e were measured and summed in each frame. After spreading, the fluorescence intensity was calculated as the mean of the values measured at three different positions inside the spread. All intensities were normalized by the total intensity inside all foci in the first scan before retraction. The data in Figure 1g (left) displaying the dependence of normalized intensity (*I*) on extension (*x*) were fit to a hyperbolic tangent function with a flat baseline on the focus side and a slanted baseline on the spread side:

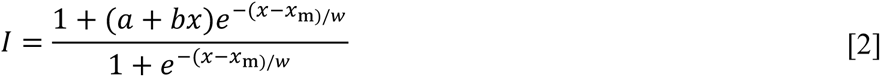

For Figure 1g (right), the total intensity over the entire spread was summed and then normalized as stated above. Only the spread immediately after the disappearance of foci was analysed, in order to minimize the effect of HMGB1 dissociation. For Figure 2d, the brightest focus in the first frame was chosen, and its intensities in successive frames were measured. After normalization by the value in the first frame, the results are displayed in Figure 2e.

### Condensate formation and imaging

HMGB1 or HMGB1-ΔC was mixed with 25-bp dsDNA or 32-nt ssDNA at final concentrations of 100 µM each in 10 mM pH 7 imidazole buffer (no KCl). Immediately after mixing them, a 2-µL aliquot was loaded onto a glass slide and imaged on an Olympus BX53 brightfield microscope using a 40× objective. Images were captured at 0 s and, for the ΔC–ssDNA sample, also at 60 s, as droplets were very crowded at 0 s. Brightfield images were similarly obtained for HMGB1 or HMGB1-ΔC homotypic condensation or their heterotypic condensation with protamine, except that the protein components were at 50 µM each and 10% PEG was present.

Conversion of protamine-dsDNA into droplets by HMGB1 was imaged on the confocal module of a LUMICKS C-Trap instrument. Aggregates were formed by mixing 9 µM protamine, 1 µM protamine-Cy5, and 20 µM 25-bp dsDNA. A 10-µL aliquot was placed in a custom sample holder built with a PEGylated coverslip attached to a glass slide using double-sided tape. Confocal scans were done using a 638 nm laser with 10% laser power. Brightfield images of the sample were also captured. In a preformed aggregate sample, 19 µM HMGB1 premixed with 1 µM GFP-HMGB1 and 10% PEG was added. Using an optical trap, droplets were trapped and moved around to fuse with other droplets, resulting in large droplets (5-10 µm in diameter) that were then parked on the PEGylated coverslip. After waiting for 5 to 10 min for any rise in temperature due to the trapping laser to dissipate, confocal scanning of droplets was performed using 638 nm (for protamine-Cy5) and 488 nm (for GFP-HMGB1) lasers at 10% laser power. This experiment was repeated with HMGB1 + GFP-HMGB1 replaced by 20 µM HMGB1-ΔC.

### Fluorescence recovery after photobleaching (FRAP)

FRAP measurements in binary and ternary droplets were performed on the LUMICKS C-Trap instrument. Binary droplets were formed by mixing 19 µM HMGB1, 1 µM GFP-HMGB1, and 20 µM of 25-bp dsDNA in 10 mM pH 7 imidazole buffer (no KCl) with 10% PEG. Ternary droplets were formed in two ways: addition of 19 µM HMGB1 + 1 µM GFP-HMGB1 and 10% PEG to aggregates preformed with 10 µM protamine and 20 µM dsDNA (same as in the preceding paragraph), or adding 10 µM protamine to an HMGB1-dsDNA binary droplet sample. Again, droplets were grown to large sizes before scanning using a 488 nm laser. The FRAP experiment on each selected droplet consisted of the following scans, with 5% laser power except for the bleaching scan, 100 nm pixel size, and 0.05 ms pixel dwell time. (1) Scan of an area covering the entire droplet. (2) Pre-bleach scan of a 0.5 µm × 0.5 µm region at the center of the droplet. (3) Bleaching scan, with 100% power for 10 s. (4) Recovery scan, for 60 s in binary droplets and 200 s ternary droplets. (5) A final full scan of the droplet. The extent of fluorescence recovery was measured from the final scan of the droplet, as the ratio of fluorescence intensity in the bleached region and the counterpart in a region away from the bleached region. The fluorescence intensity in the bleached region as a function of post-bleach time was fit to

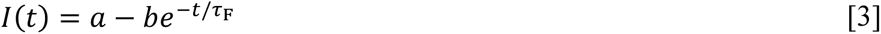

The fit was limited to the first 50 s in order to minimize the effect of photobleaching during the recovery scan. Five replicates were performed for each type of droplets.

### OT-directed droplet fusion

Binary and ternary droplets were prepared similarly to the FRAP experiment. HMGB1 and 25-bp dsDNA (each at 20 µM final concentration) were mixed in 10 mM pH 7 imidazole buffer (no KCl) with 10% PEG to produce binary droplets. Addition of 10 µM protamine to binary droplets produced ternary droplets. The two optical traps of the C-Trap instrument (50:50 split of laser power) were used to trap two droplets and grow to an equal size. The overall laser power was then reduced to 3%. With the brightfield camera on, the trap-1 droplet was moved slowly to be in contact with the trap-2 droplet. Fusion then proceeded spontaneously. The force on trap 2 was recorded at a sampling rate of 78.125 kHz. The time course of the normalized trap-2 force was fit to^53^

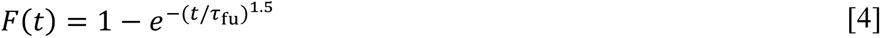

to obtain the fusion time 𝜏_fu_. The brightfield video was analyzed using ImageJ to determine the pre- and post-fusion droplet radii.

### Salt dissolution of HMGB1-DNA droplets

20 µM HMGB1-WT or HMGB1-ΔC was mixed with 20 µM 25-bp dsDNA or 31.25 µM 32-nt ssDNA (for charge matching between dsDNA and ssDNA) in 10 mM pH 7 imidazole buffer with 0 to 1 M KCl and containing 10% PEG. A 2-µL aliquot was loaded onto the lower pedestal of a NanoDrop 2000c spectrophotometer and the absorbance at 350 nm was measured. Four replicates were performed at each salt concentration. The mean absorbance (*A*_350_) as a function of salt concentration ([KCl]) was fit to

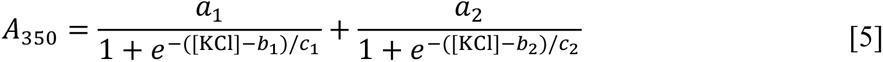

The concentration at which *A*_350_ decayed to the background level, 0.005, was taken as the dissolution [KCl].

## Acknowledgement

We thank Dr. Geeta Narlikar for gifting us the plasmids of HMGB1 and HMGB1-ΔC. This work was supported by National Institutes of Health Grant GM118091.

