## Supplementary Table and Figures for "Counteraction of HMGB1 at ss-dsDNA junctions maintains liquidity of protamine-DNA co-condensates"

*Supporting Information*

Table S1. Sequences and domain features of sperm nuclear basic proteins in flies.

| Protein[a] | Sequence and annotation[b] | #R/K<br>#D/E[c] |
|---|---|---|
| Transition protein-like | | |
| Tpl94D<br>(Q8IMZ6) | MGSVLSRSDQPRDSVAYRNFLTVYRNQHKQMSSNDALLSASREWGTLSQAQRNRYANMKDSPKRGTKNITENAAA**RRKITKKNSRSKRISQRR**IQDKGSAYKPLTLNRSYVIRKSLVPSRGFLNLVQFFQEIHSEMPRSLAVKEAVSSWCAMNADQRGIFISDL | 10 |
| tHMG1<br>(Q9VCZ5) | MSDCGARPKKPMSAFMLWMNSTGRKHIKAEHPDFSVQEVSVKGGEMWRAMADEDKIVWQESATTAMAEYKEKLKQWNFPKEHRFSDTQCICSSNTNQCPTLFVYDTMDDSMTPIC**RKCLSKTR**CLH | 4 |
| tHMG2<br>(Q9VCZ4) | MSDYGARPKKPMSAFMLWMNSTGRKNIRAEHPDFSVQEVSVKGGEMWRAMADEHKIVWQESASKAMAEYKEKLEKWNAFKEHQTESFPHIYEAPLSSRFSKTNQRPTLFVYDSKDEAMAPIC**RTCFSKAK**CFH | 3 |
| CG30356<br>(A1Z7I6) | MGRAVKNNRVRSAHCNSAAKATSISGHSMLKQNNDCNGRNPFFQFLAYFRKCSNNCLGHLPSDRVTLIAGKVWNYMSLSEKEPFIAAARRFNYTYRS**RSRKVNWVLAQLRK**SAAGEECRPQAQWMLMNFLKSWQESVVRNLLDLDHNQN | 5 |
| Protamine-like | | |
| ProtA<br>(Q9V3S3) | MSSNNVNECKSLWNGIISISAKDESPKGLTEMCNHPI**RRAPQKCKPMKSCAKPRRKAACAKATRPKVKCAPRQKCSK**QGPVTNNAYLNFVRSFRKKHCNLKPRELIAKAAKAWARLSENRKDRYRRMACKVTTSERHKRRRICQQY | 16 |
| ProtB<br>(Q9VJR1) | MSSNNVNECKSLWNGIISISAKDESPKGLTEMCNHP**KRRAPPKCKPMKSCAKPRRKAACAKATRPKVKCAPSQKCSK**QGPVTNNAYLNFVRFFRKKHCDLKPQELIAEAAKAWAELPEHRKDRYRRMACKVTTSERHKRRRICK | 16 |
| Mst77F<br>(P16909) | MSNLKQKDSKPEVAVTKSVKTYKKSIEYVNSDASDIEEDINRAEDEYASSSGFVNFLRDFKKRYGEYYSNNEIRRAAETRWNEMSFRHRCQYSAEPLDTFHVEPNSVSSLQRSSEGEHRMHSEISGCADTFFGAGGSNSCTPRKEN**KCSKPRVRKSCPKPRAKTSKQRRSCGKPKPKGARPRKACPRPRKK**MECGKAKAKPRCLKPKSSKPKCSM | 21 |
| Prt99C<br>(Q9VAF7) | MGRKRGRKEYCPPIYKRQKVARVTNNGYLNFMTEYKKRFYGLSPQDMVHYAAKQWTQLSMAEKEAFKSKKPSTITLKSPAQYVACEMKSDVAGGQQSSCQRQSPSARLRESE**RRSSRSKTLCRSAKNRQRGKPKPQQSKRR**LSHMGSAVAYIHFLRKFQRKNTELRTIDLLKTATRLWCRLPERHRHAFERPLWIVTIGKS | 13 |
| CG14835<br>(Q9VS70) | MDYSYSIGSKSPVFISKCAPYNFLREYRLGCDCKNIPAVTLVKEARNAWHALSDQEQSLFEEVPYLMAQFGPSIESAMRLTVNVLSPQQVTVQRTRSDVQSS**KRRICRGNAKTHRKKRQQKMGATSRKR**SPGIGSTRCTRYTSQFHWENPPL | 13 |
| ddbt<br>(A2VEN4) | MARSQSKMIRAAMVVDDPNMVAWLPYLNFLRFLKRNFYPRTDLRRLLQVGLIRWIALSDAQKRLFEPERILA**RVARRQRNKRRRRLLRRARHGQKGRGAVRRPIYDSRPKPRRRKPK** | 23<br>1 |

[a]Uniprot ID is given in parentheses after the protein name.

[b]HMG boxes are underlined while Arg/Lys-rich motifs are in bold. HMG boxes were determined based on InterPro (https://www.ebi.ac.uk/interpro/search/sequence/), Chang et al. (eLife 2023), and the consensus F(L/M/V)…Wxxxxxxxxxx(F/Y/W). Consensus residues are shaded in green.

[c]Number of Arg and Lys residues (and number of Asp and Glu residues if any) in Arg/Lys-rich motifs, identified as the shortest stretch of sequence outside HMG boxes with the highest net charge and at most one D or E.

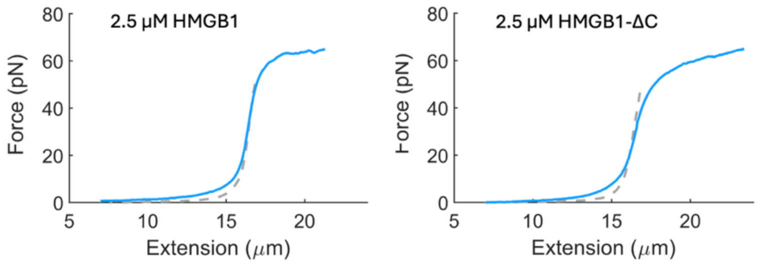


Figure S1. Stretching curves of λ-DNA in the presence of 2.5 μM HMGB1 or 2.5 μM HMGB1-ΔC.

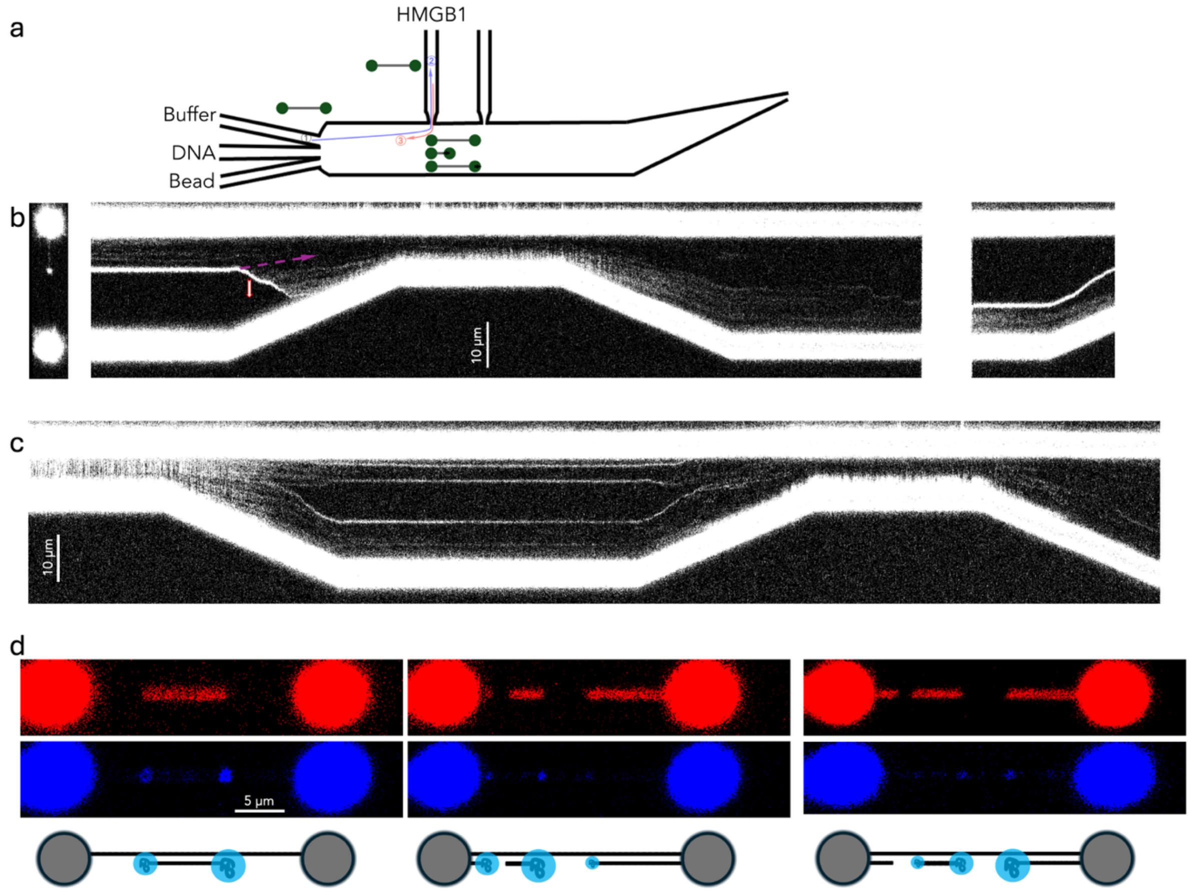


Figure S2. HMGB1 focus formation and localization on λ-DNA. (**a**) General protocol for the studies in the presence of HMGB1 alone. A λ-DNA tether was first overstretched in the buffer channel, then soaked in the HMGB1 channel, and finally brought outside the channel for retraction-stretching cycles and confocal scanning. (**b**) A kymograph, similar to Figure 1h, showing a focus spreading on overstretched λ-DNA upon retraction. A slanted arrow indicates the would-be direction of a static point during retraction; a vertical arrow indicates the migration direction of a focus. A 2D scan before retraction is displayed at the left; a kymograph after dipping the re-stretched tether in the HMGB1 channel is displayed at the right. (**c**) A kymograph following an alternative protocol, where a λ-DNA tether started in a relaxed state (extension at 7 μm) underwent stretch-retract cycles. In (**b**) and (**c**), the channel contained 8 μM HMGB1 and 0.8 μM GFP-HMGB1. (**d**) Top: YOYO-3 fluorescence of λ-DNA tethers at overstretched to 23, 21, and 20 μm, respectively. Middle: GFP-HMGB1 fluorescence, showing foci at ss-dsDNA junctions. The channel contained 5 μM YOYO-3 and 2.5 μM GFP-HMGB1; scanning took place immediately after the tethers were moved outside the channel. Bottom: carton illustration.

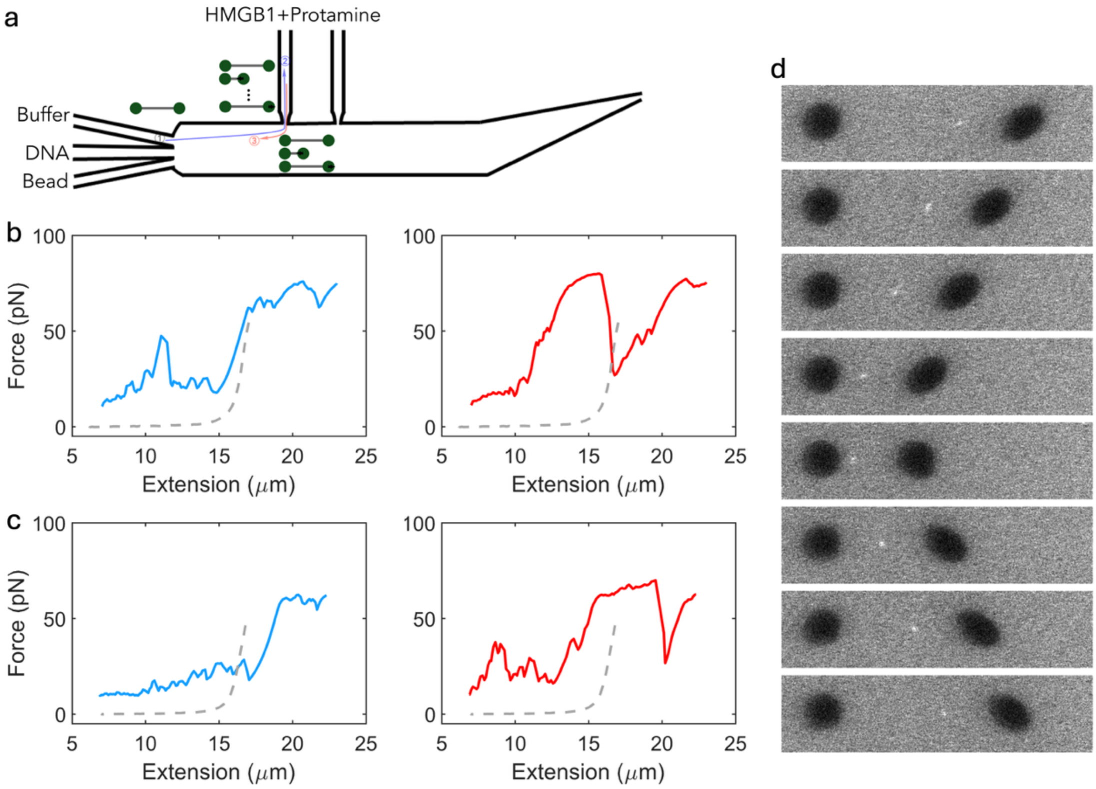


Figure S3. HMGB1's counteraction to protamine. (**a**) General protocol for the studies in the simultaneous presence of HMGB1 and protamine. A λ-DNA tether was first overstretched in the buffer channel, then transferred to the HMGB1 + protamine channel for soaking or multiple retract-stretch cycles, and finally brought outside the channel for one more retraction-stretching cycle and confocal scanning. (**b, c**) Same as Figure 2b, with force-extension curves from two more tethers. For each tether, the curve (blue color) in the left panel was acquired when the tether was inside the HMGB1 + protamine channel (at 5 and 2.5 μM, respectively), the curve (red color) in the right panel was acquired after the tether was moved just outside the channel. (**d**) 2D scans of a λ-DNA tether undergoing retraction and stretching inside the HMGB1 + protamine channel (8 μM HMGB1; 0.8 μM HMGB1; and 2.5 μM protamine). Fluorescence was rendered black and white.

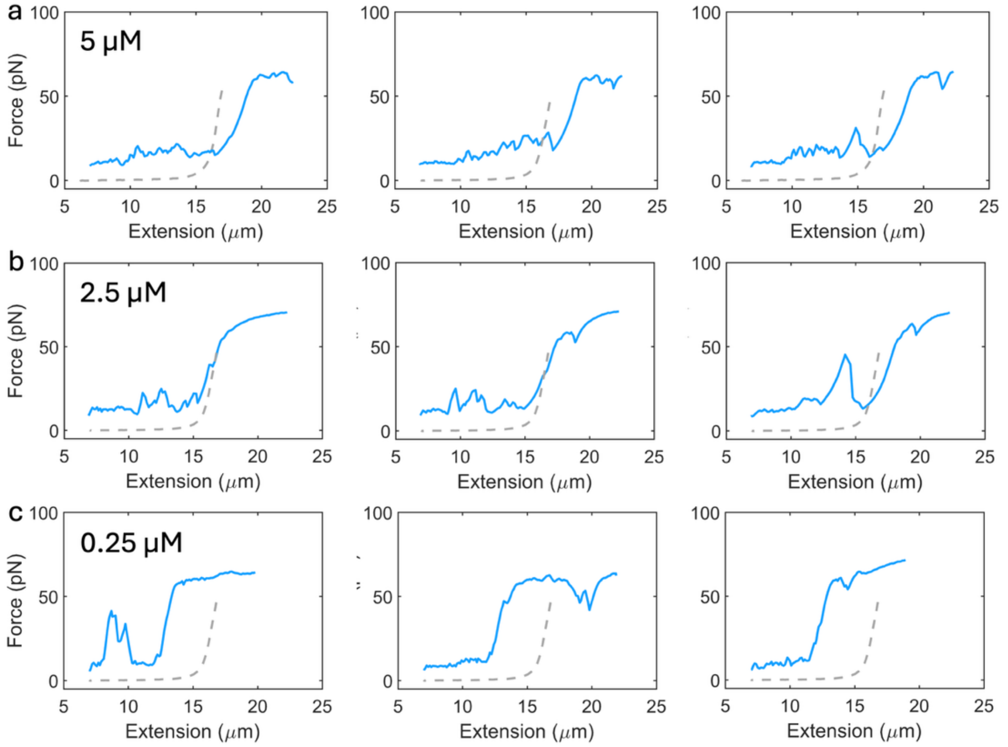


Figure S4. Force-extension curves of λ-DNA during consecutive retract-stretch cycles in the HMGB1 + protamine channel. (**a-c**) HMGB1 at 5, 2.5, and 0.25 μM, respectively; protamine was fixed at 2.5 μM. Each row presents the first three consecutive stretching curves.

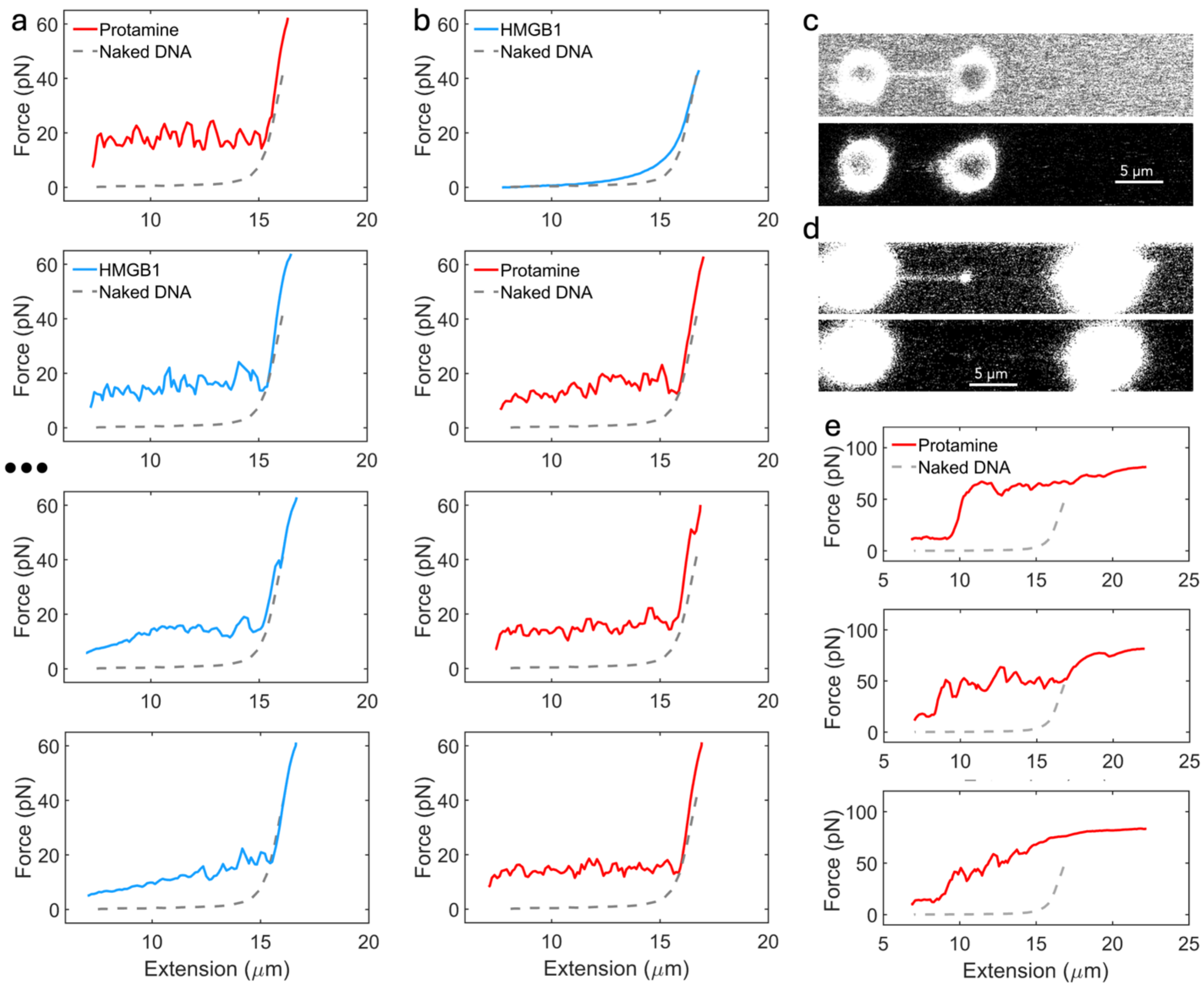


Figure S5. Effects of protamine and HMGB1, when placed in different channels, on λ-DNA tethers. (**a**) Consecutive stretching curves of an intact-DNA tether, first placed in protamine (2.5 μM; red curve) and then moved into HMGB1 (2.5 μM; blue curves). Three stretching curves are skipped (indicated by ellipses); subsequently, the initial values of the force curves shifted toward 0, suggesting some protamine dissociation in the HMGB1 channel. (**b**) Consecutive stretching curves of an intact-DNA tether, first placed in HMGB1 (2.5 μM; blue curve) and then moved into protamine (2.5 μM; red curves). (**c**) Top: a 2D scan when a relaxed tether (7 μm) was inside the HMGB1 channel (8 μM HMGB1 with 0.8 μM GFP-HMGB1), showing a uniform layer of HMGB1 spread over the DNA surface. Bottom: a 2D scan after the tether was moved into protamine (2.5 μM), showing the absence of an HMGB1 layer. (**d**) Similar to (**c**), but the tether was initially overstretched (to 23 μm). Top: scan taken immediately after the tether was moved outside the HMGB1 channel, showing a focus with strong fluorescence; bottom: scan taken after the tether was then moved to the protamine channel, showing a very faint focus. (**e**) Consecutive stretching curves inside protamine (2.5 μM), after an initially overstretched tether was soaked in the HMGB1 channel (8 μM HMGB1 with 0.8 μM GFP-HMGB1).

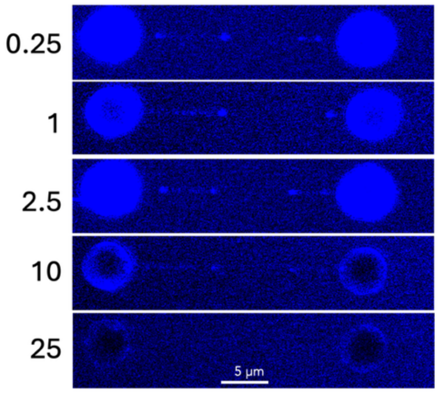


Figure S6. Displacement of GFP-HMGB1 from foci by HMGB1. Shown are 2D scans of overstretched λ-DNA tethers in the presence of 0.25 µM GFP-HMGB1 and HMGB1 at the indicated concentrations in µM. For each tether, the second of three consecutive scans is displayed.

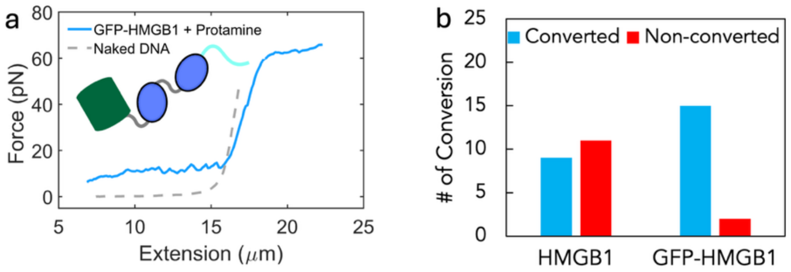


Figure S7. GFP-HMGB1 was more effective than HMGB1 in tangle-to-bridge conversion. (**a**) A force-extension curve showing conversion of tangles to bridges by 2.5 μM GFP-HMGB1. (**b**) A bar graph showing the number of λ-DNA tethers for which tangles were or were not converted into bridges in the (GFP-)HMGB1 + protamine channel (2.5 μM protamine along with 2.5 μM HMGB1 or GFP-HMGB1). The HMGB1 results are the same as in Figure 2c.

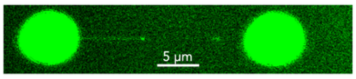


Figure S8. A 2D scan of an overstretched λ-DNA tether (extension ~ 23 µm) in a channel containing 8 µM HMGB1 and 0.8 µM HMGB1-Cy3. Two foci were formed, along with a faint coating of the dsDNA sections to their left.

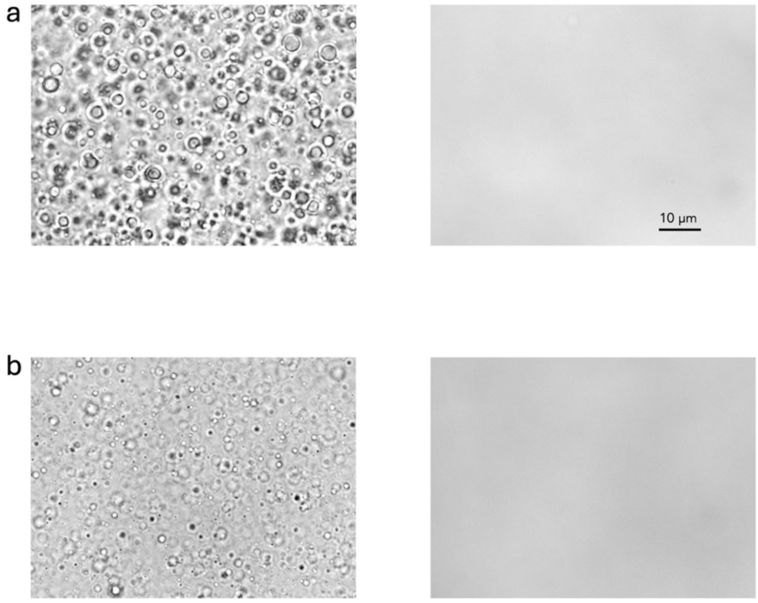


Figure S9. Effects of the acidic tail on the homotypic condensation of HMGB1 and the heterotypic condensation of HMGB1 and protamine. (**a**) Brightfield images showing droplet formation of HMGB1 (left) and absence of condensation by HMGB1-ΔC (right). (**b**) Brightfield images showing droplet formation of a HMGB1−protamine mixture (left) and absence of co-condensation by HMGB1-ΔC and protamine (right). Protein components were at 50 µM each and dissolved 10 mM imidazole buffer (pH 7) with 10% PEG.

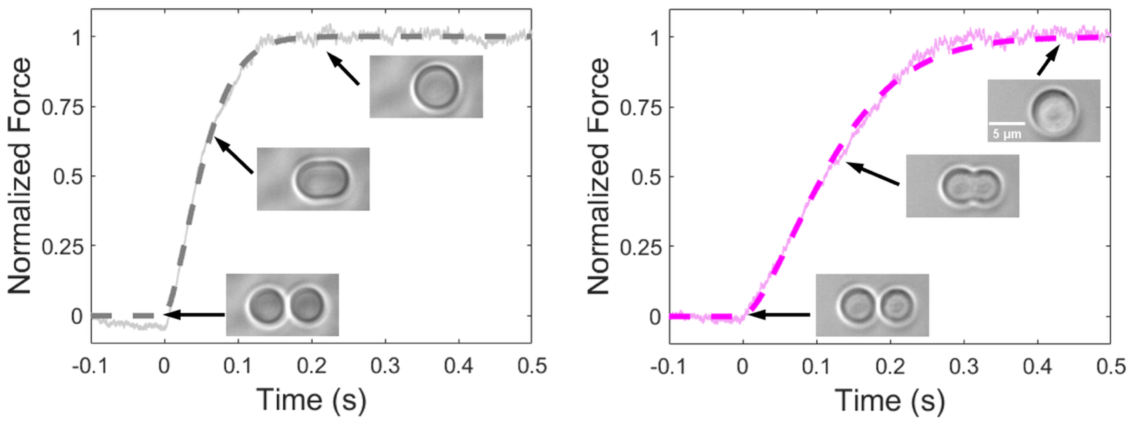


Figure S10. Protamine slowed down the fusion of HMGB1–dsDNA droplets. (**a**) The fusion progress curve of a pair of equal-sized HMGB1–dsDNA droplets. Droplets were formed by mixing 20 µM HMGB1 with 20 µM 25-bp dsDNA in 10 mM imidazole (pH 7) with 10% PEG and grown to ~2.5 µm radius before fusion. (**b**) Corresponding results for HMGB1–dsDNA–protamine droplets. Droplets were prepared by adding 10 µM protamine to the sample in (**a**). Solid traces display raw data; dashed curves are fits to equation [4].
